# RESCRIPt: Reproducible sequence taxonomy reference database management for the masses

**DOI:** 10.1101/2020.10.05.326504

**Authors:** Michael S. Robeson, Devon R. O’Rourke, Benjamin D. Kaehler, Michal Ziemski, Matthew R. Dillon, Jeffrey T. Foster, Nicholas A. Bokulich

## Abstract

**Background:** Nucleotide sequence and taxonomy reference databases are critical resources for widespread applications including marker-gene and metagenome sequencing for microbiome analysis, diet metabarcoding, and environmental DNA (eDNA) surveys. Reproducibly generating, managing, using, and evaluating nucleotide sequence and taxonomy reference databases creates a significant bottleneck for researchers aiming to generate custom sequence databases. Furthermore, database composition drastically influences results, and lack of standardizations limits cross-study comparisons. To address these challenges, we developed RESCRIPt, a software package for reproducible generation and management of reference sequence taxonomy databases, including dedicated functions that streamline creating databases from popular sources, and functions for evaluating, comparing, and interactively exploring qualitative and quantitative characteristics across reference databases.

**Results:** To highlight the breadth and capabilities of RESCRIPt, we provide several examples for working with popular databases for microbiome profiling (SILVA, Greengenes, NCBI-RefSeq, GTDB), eDNA, and diet metabarcoding surveys (BOLD, GenBank), as well as for genome comparison. We show that bigger is not always better, and reference databases with standardized taxonomies and those that focus on type strains have quantitative advantages, though may not be appropriate for all use cases. Most databases appear to benefit from some curation (quality filtering), though sequence clustering appears detrimental to database quality. Finally, we demonstrate the breadth and extensibility of RESCRIPt for reproducible workflows with a comparison of global hepatitis genomes.

**Conclusions:** RESCRIPt provides tools to democratize the process of reference database acquisition and management, enabling researchers to reproducibly and transparently create reference materials for diverse research applications. RESCRIPt is released under a permissive BSD-3 license at https://github.com/bokulich-lab/RESCRIPt.

## Background

Marker-gene amplicon and metagenome sequencing have become attractive methods for characterizing microbial community composition and function [1,2] in human health [3–5] and agriculture [6–8], as well as macroorganism diversity through diet metabarcoding studies [9–11] and environmental DNA (eDNA) surveys [12–15]. Taxonomic classification is often a primary goal in marker-gene and metagenome sequencing studies to identify the composition of a mixed community, or to detect species of interest (e.g., pathogens). This is accomplished by comparing the observed sequences to a reference database consisting of target marker-gene or genome sequences from known species. The selection of a reference database can significantly impact both marker-gene and metagenome sequencing results [16,17], and methods for assessing database quality and fitness for a given sample type or hypothesis remain an undermet need.

Identification of Bacteria and Archaea is most commonly performed using the 16S rRNA gene, due to its historical use as a phylogenetic marker [18,19] and the existence of curated reference databases [20,21]. The SILVA [20,22] rRNA gene database and Greengenes [21] 16S rRNA gene databases are commonly used for identifying Bacteria and Archaea, containing curated taxonomies, sequences, and phylogenies. More recently, the Genome Taxonomy Database (GTDB) was developed with the intent to provide a standardized bacterial and archaeal taxonomy based on genome phylogeny [23,24], and provides 16S rRNA reference sequences. NCBI-RefSeq also provides several targeted loci sequence databases from curated records, including Internal Transcribed Spacer (ITS), and both the small and large sub-unit (SSU & LSU) rRNA genes [25]. Non-16S genes are also attractive targets for bacterial and archaeal species identification due to the degree of species resolution that they afford, but their application is limited by the relative lack of curated reference materials [26–28].

Fungal classification is most commonly performed using the ITS domain, the designated fungal “barcode of life”, though the SSU and LSU rRNA genes are also common targets [29]. Both NCBI-RefSeq [25] and the UNITE database [30] provide curated ITS sequences from fungi and other eukaryotes, as well as the RDP Warcup fungal ITS training set [31], which was prepared from an earlier release of the UNITE+INSD database. Both SILVA [22] and RDP [32] provide LSU databases for fungal sequence classification. NCBI RefSeq releases databases for both fungal SSU and LSU [25].

Macroorganism identification, for both diet metabarcoding and eDNA surveys, is commonly accomplished using the mitochondrial cytochrome oxidase subunit 1 (COI) gene for metazoa [33–35], ITS2 and chloroplast *trn*L (UAA) intron [36–38] for plants, 12S rRNA for fish [39,40], and a variety of other clade-specific marker genes. For some of these marker genes, curated reference databases exist, such as BOLD for COI [33] and PLANiTS for plant ITS2 [38], but for others the process of generating custom reference databases poses a research bottleneck.

Taxonomic profiling studies rely on high-quality sequence taxonomy reference databases. However, errors in public sequence databases are well documented [12,41,42] and can lead to misclassification errors in downstream results [12]. Different reference databases can yield widely different classification results for biological data, but standards are lacking to objectively assess the quality of individual databases [43]. Revisions to taxonomic naming [44–49] and the rapid pace at which new sequences and genomes are added to public databases mean that curated reference releases may lag behind [50]. Additionally, issues with amplicon length and sequence heterogeneity can limit the ability to identify species, especially from short marker-gene sequences or metagenome fragments [51]. Hence, many researchers choose to perform additional curation to focus on type strains [52], quality filtering [14,53], or construct environment-specific databases that are constrained to contain species found within a given environment [52,54–59]. Database customization is also often performed to add new accessions that are absent in some database releases to increase database coverage [50], or to incorporate outgroups [14]. However, generating such databases can be technically challenging, subjective, and difficult to document, leading to issues with transparency and reproducibility, and limiting the ability of many researchers to acquire appropriate reference materials for their studies, or leading to reliance on proprietary resources and services (limiting scientific transparency and increasing research costs). Sequence curation is a significant hurdle in this process, as taxonomic misannotations, sequence errors, and other errors in existing (and inchoate) reference sequence databases reduce the accuracy of taxonomic classifiers that rely on these data [41,42,60–62].

The need for scientific results to be reproducible, replicable, and transparent has taken on new urgency in the digital age [63]. On the one hand, increasing experimental and analytical complexity pose mounting challenges to effective documentation and sharing of methodological procedures and results [63–66]. On the other hand, digital tools present opportunities to address these challenges, and various reporting standards have been published to guide researchers in reporting and publishing new types of data, software, and other resources [67–69]. Following guidelines such as these is important for reporting, but also for standardization of methods during data reuse and metaanalysis. Given the fundamental importance of reference databases to reporting results from marker-gene and metagenome experiments, it is critical that principles such as these be followed by researchers when they acquire, modify, and use reference data.

To address the need for reproducible bioinformatics workflows to streamline database generation and curation, we developed RESCRIPt (REference Sequence annotation and CuRatIon Pipeline; https://github.com/bokulich-lab/RESCRIPt), a Python 3 package for retrieving, filtering, and evaluating nucleotide sequence and taxonomic data (Figure 1). RESCRIPt was implemented as a QIIME 2 [70] plugin, in order to incorporate the integrated data provenance and multiple user interfaces of QIIME 2. Hence all processing steps used to generate a database are recorded in provenance that is stored both in the database files as well as in all downstream results, enhancing scientific transparency, reproducibility of results, and replication of the database (and the processing steps used in its creation) by other researchers, following the FAIR data principles to make data findable, accessible, interoperable and reusable [67]. RESCRIPt enables efficient and transparent construction of reference databases for any amplicon targets for which source data exist. It also enables researchers to evaluate the information content of sequence taxonomy databases, including sequence and taxonomic entropy, unique labels, and taxonomic classification accuracy. We used RESCRIPt to benchmark several commonly used reference sequence databases side-by-side to evaluate relative information and performance characteristics, before and after various curation steps. Finally, we demonstrate the broad applications of RESCRIPt for creating reproducible and extensible sequence analysis workflows via a comparison of hepatitis genomes from several global sources.

**Figure 1.**
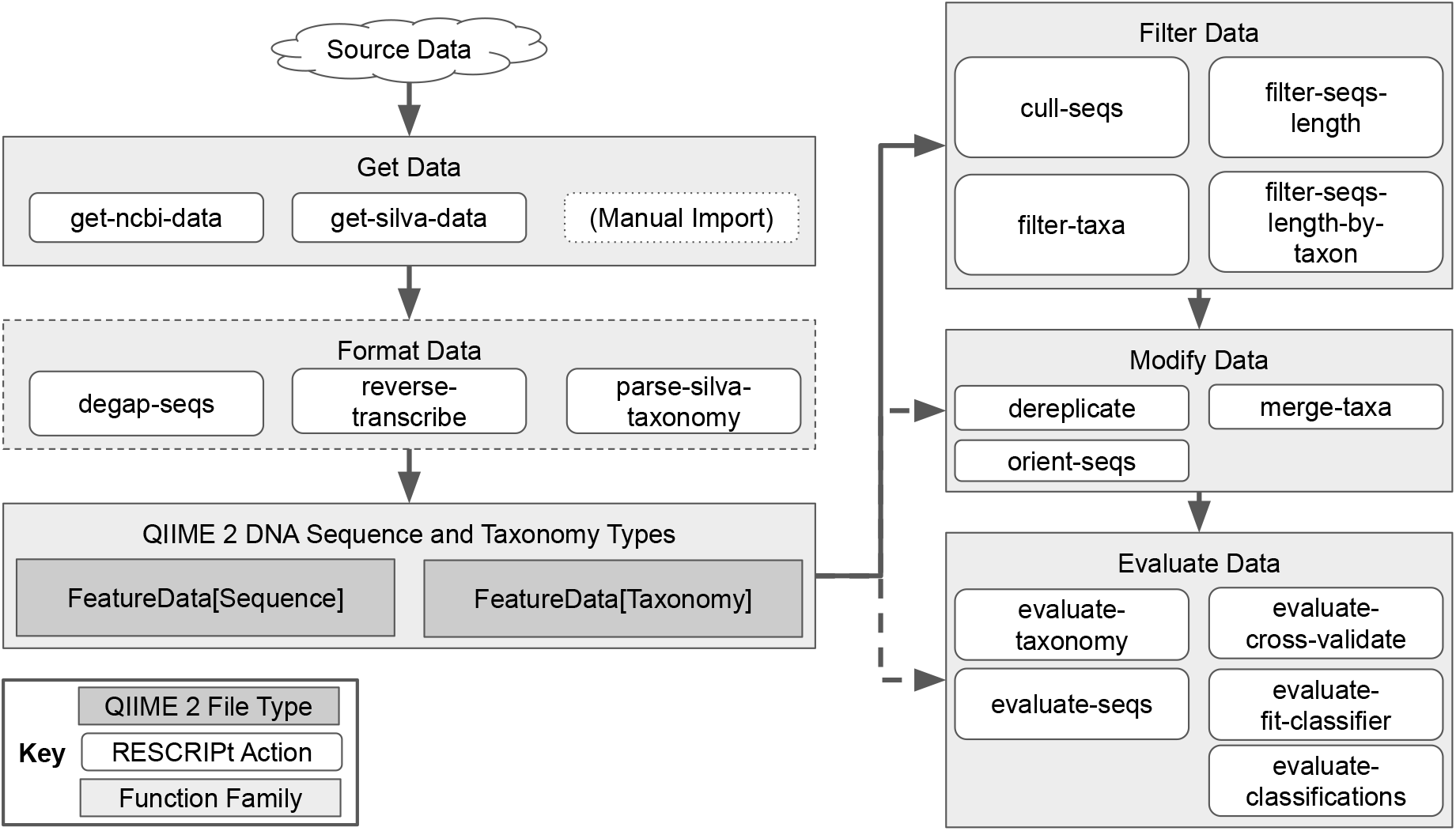
Current RESCRIPt functionality for processing and curating reference sequence data. Arrows indicate suggested workflows. Dotted arrows and edges indicate optional steps for customized workflows.

## Results

### Comparison of 16S rRNA Gene Sequence Databases

Researchers investigating bacterial and archaeal community compositions using 16S rRNA gene sequences are faced with myriad options for reference sequence databases. Those using non-16S genes will be quick to remind them that having choices is a good problem to have — but selecting the “best” reference materials for a specific task is still indeed a problem faced by many researchers. Although the Greengenes database [21] was popular among the microbiome research community in the initial years after its first release, its popularity has waned in recent years as (at the time of writing) the last release was in 2013 and much has changed in the world of microbial taxonomy in the interim. The SILVA database [22] has been a popular alternative, boasting a regular release cycle, curated taxonomy and sequences, and a large database size. More recently, the GTDB project [23] seeks to create a standardized bacterial and archaeal taxonomy based on genome phylogeny, making it an attractive database for some researchers. Meanwhile, many other options exist, such as NCBI-RefSeq [25] for a curated set of type strains and high-quality reference genomes. We conducted a benchmark of these four 16S rRNA gene databases using RESCRIPt to compare various qualitative and quantitative characteristics and performance metrics. This benchmark was performed using full-length 16S rRNA gene sequences from each database, so the goal was to compare full-length sequence and taxonomy information, not to simulate performance for commonly used short-read sequencing technologies.

RESCRIPt’s evaluate-seqs action was used to examine sequence length distributions (Figure 2A), the number of unique sequences (Figure 2B), and sequence and kmer entropy as measures of both richness and evenness of unique sequences in these databases (Figure 2C). The evaluate-taxonomy action was used to examine the number of unique taxonomic labels (Figure 3A), taxonomic entropy (Figure 3B), and the number of unclassified labels at each taxonomic rank (Figure 3C). The evaluate-fit-classifier and evaluate-classifications actions were used to compare optimal classification performance for each classifier (Figure 3D).

**Figure 2.**
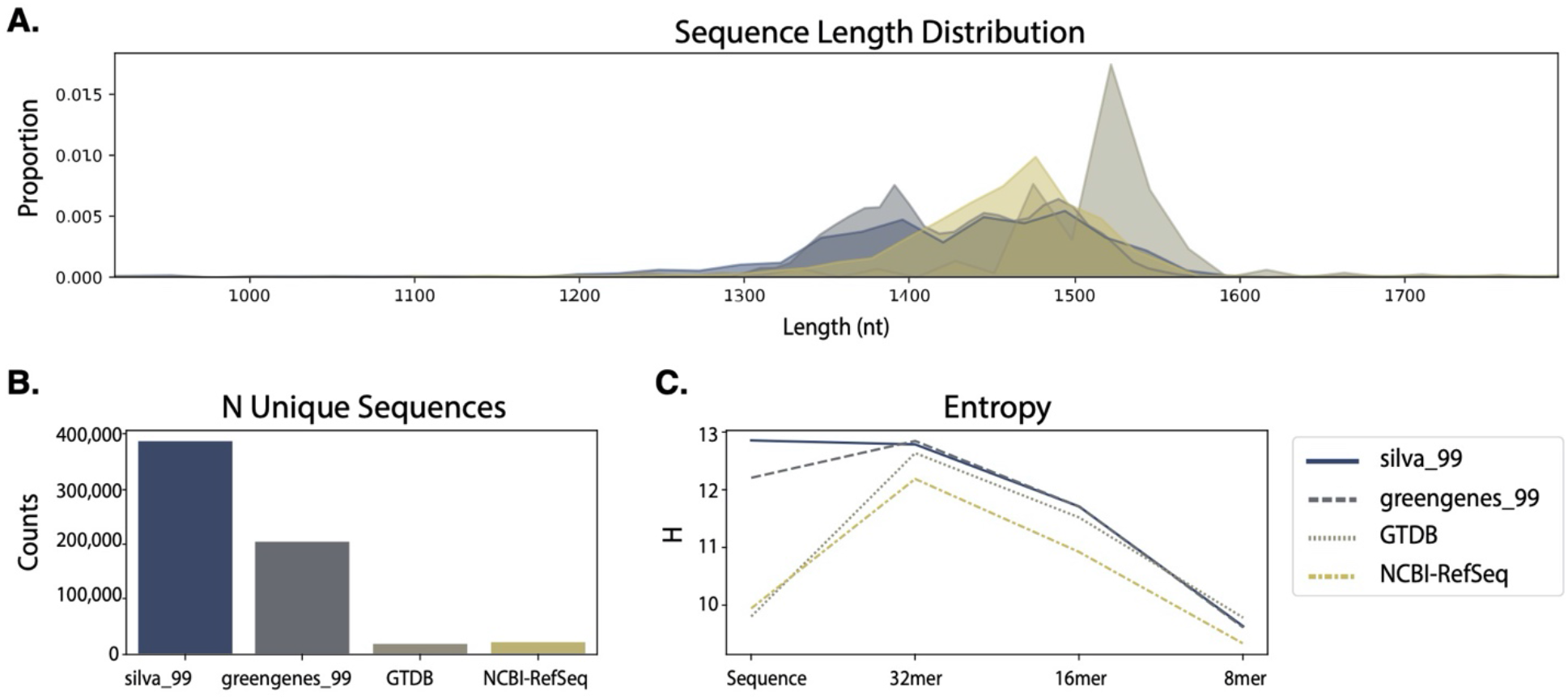
Comparison of sequence information from SILVA, Greengenes, GTDB, and NCBI-RefSeq 16S rRNA gene databases. A, Sequence length distributions (after removing outliers, see materials and methods). B, Number of unique sequences in each database. C, Entropy of full-length sequences and different kmer lengths in each database.

**Figure 3.**
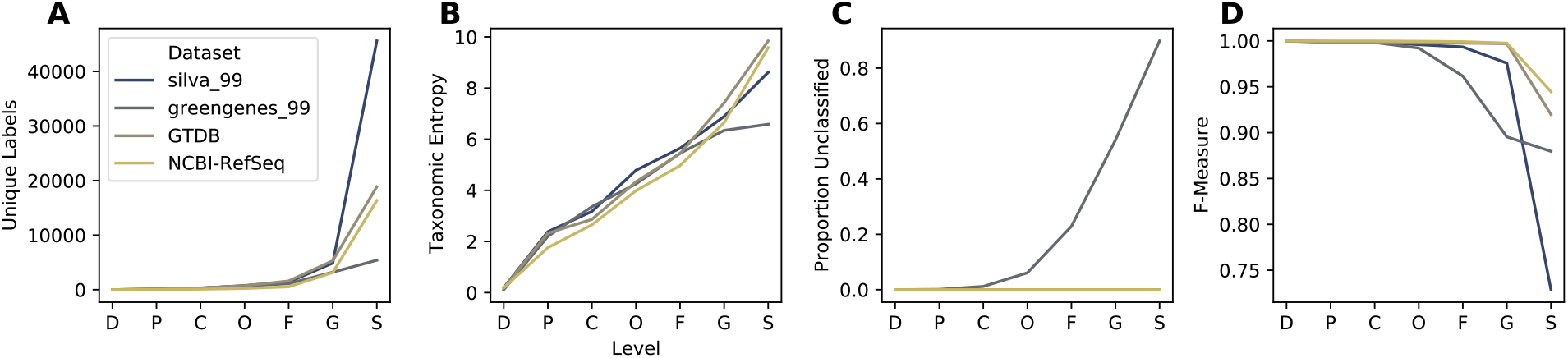
Comparison of taxonomic information and simulated classification accuracy from SILVA, Greengenes, GTDB, and NCBI-RefSeq 16S rRNA gene databases. A, Number of unique taxonomic labels; B, Taxonomic entropy; C, proportion of unclassified taxa at each rank; D, optimal classification accuracy (as F-Measure) without cross-validation (simulating best possible classification accuracy when the true label is known but classification accuracy may be confounded by other similar hits in the database). Cross-validation was not used because two of the databases (GTDB and NCBI-RefSeq) lack replicate species. Rank labels on x-axis: D = Domain, P = phylum, C = class, O = order, F = family, G = genus, S = species.

Results illustrate varying length distributions across databases, reflecting different proportions of Bacteria and Archaea in each, as well as different methods used to identify start and end sites in each of these databases. Notably, some outliers (as short as 200 nt and as long as 3983 nt) were initially observed in SILVA, NCBI, and GTDB, presumably representing partial and untrimmed 16S rRNA gene sequences (data not shown). These were removed prior to downstream evaluation to avoid biasing performance metrics (see Methods section). Researchers should be aware of length aberrations in these and other reference databases, and can use the evaluate-seqs action in RESCRIPt to check length distributions in their own databases before proceeding.

The SILVA database exhibited the highest number of unique sequences (Figure 2B) and species labels among the databases compared here (Figure 3A). However, only ∼25% of the species annotations used by SILVA are useful and match the genus annotations. Among these, 72% of species labels consist of unidentified, uncultured, or unknown organisms, and 2.5% of the remaining labels (excluding chloroplast and mitochondrial sequences) do not match the genus (data not shown). Notably, this is because SILVA only curates the taxonomy to genus level but provides the “organism name” given to the sequence in the NCBI GenBank source data, and hence genus–species mismatches are not uncommon. Furthermore, GTDB and Greengenes display similar levels of kmer entropy (Figure 2C), suggesting that these databases cover a similar sequence space and taxonomic diversity to SILVA. GTDB actually has more species-level annotations than SILVA (once unidentified and mismatched species labels are discounted), and higher species label entropy (Figure 3B), indicating less redundancy. The lack of species-rank curation in SILVA leads to poor optimal classification performance at the species level (Figure 3D), yielding a species-level F-measure of 0.73, far below the other 16S rRNA gene databases. By comparison, classification accuracy at the genus level is much higher for SILVA, consistent with the level of curation performed (Figure 3D)

NCBI-RefSeq and GTDB share the highest species-level taxonomic entropy, and the most unique species, after discounting the unknown/unmatched labels in SILVA (Figure 3A-B). However, NCBI-RefSeq has fewer unique sequences and lower sequence entropy (Figure 2B-C), indicating that a lower amount of sequence space is covered, most likely because of the stringent quality control and assessment process employed. Thus, NCBI-RefSeq exhibits high quality, but may have limited coverage for characterization of microbial communities in some environments, e.g., where a large number of unknown species may be encountered. In well characterized environments, this database is likely to offer competitive advantages in terms of its curated taxonomy, size, and extensive use of type strains. NCBI-RefSeq exhibited the highest optimal classification accuracy (F = 0.94, Figure 3D), though this is likely aided by the smaller database size and use many genomes sequenced from type material, reducing taxonomic ambiguities that are likely to occur in nature and are reflected in the other databases. GTDB exhibited a slightly lower optimal classification accuracy (F = 0.92), indicating very high optimal accuracy in spite of its size, suggesting that the curation efforts and taxonomic re-classification strategies employed by the GTDB curators lead to a very well resolved (if currently not officially recognized) taxonomic labeling scheme that closely aligns with the 16S rRNA gene sequence space.

Greengenes 13_8 hosts a large number of unique sequences (Figure 2B) and similar sequence entropy to SILVA (Figure 2C), but the use of 99% OTU clustering and LCA for assigning taxonomic labels to sequence clusters yields many sequences that are unannotated at the genus (54%) and species (90%). This indicates that a large number of sequences in this database are genetically similar (≥ 98%) but taxonomically distinct, yielding ambiguous labels. This highlights practical disadvantages with using this database, as 99% OTU clustering (the highest % similarity provided in the 13_8 release) limits sequence information that could be used to differentiate groups (e.g., dereplication to 100% OTUs would preserve this sequence diversity). In practice, the use of non-sequence information (e.g., ecological distribution) can be leveraged to guide taxonomic classification [54] and hence preserving this information can be advantageous in some use cases. RESCRIPt provides users with a variety of taxonomy dereplication options to put such decisions in the hands of individual researchers, and to make data processing pipelines transparent and reproducible so that others can reconstitute and adjust processing decisions as desired.

To provide context to database size comparisons, we evaluated taxonomic overlap among these databases, extending an earlier comparison of SILVA, Greengenes, NCBI (not RefSeq), and other taxonomies by Balvociute and Huson [71]. Figure 4 illustrates the proportion of labels shared at each taxonomic rank, between each pair, trio, and across all databases, relative to the total number of taxonomic labels in each database. Labels that were a prefix of another label were collapsed into that label to avoid undercounting the number of shared labels between databases, and to account for subclade labels used by GTDB (for example, “Lactobacillus_A” would be considered the same label as “Lactobacillus”).

**Figure 4.**
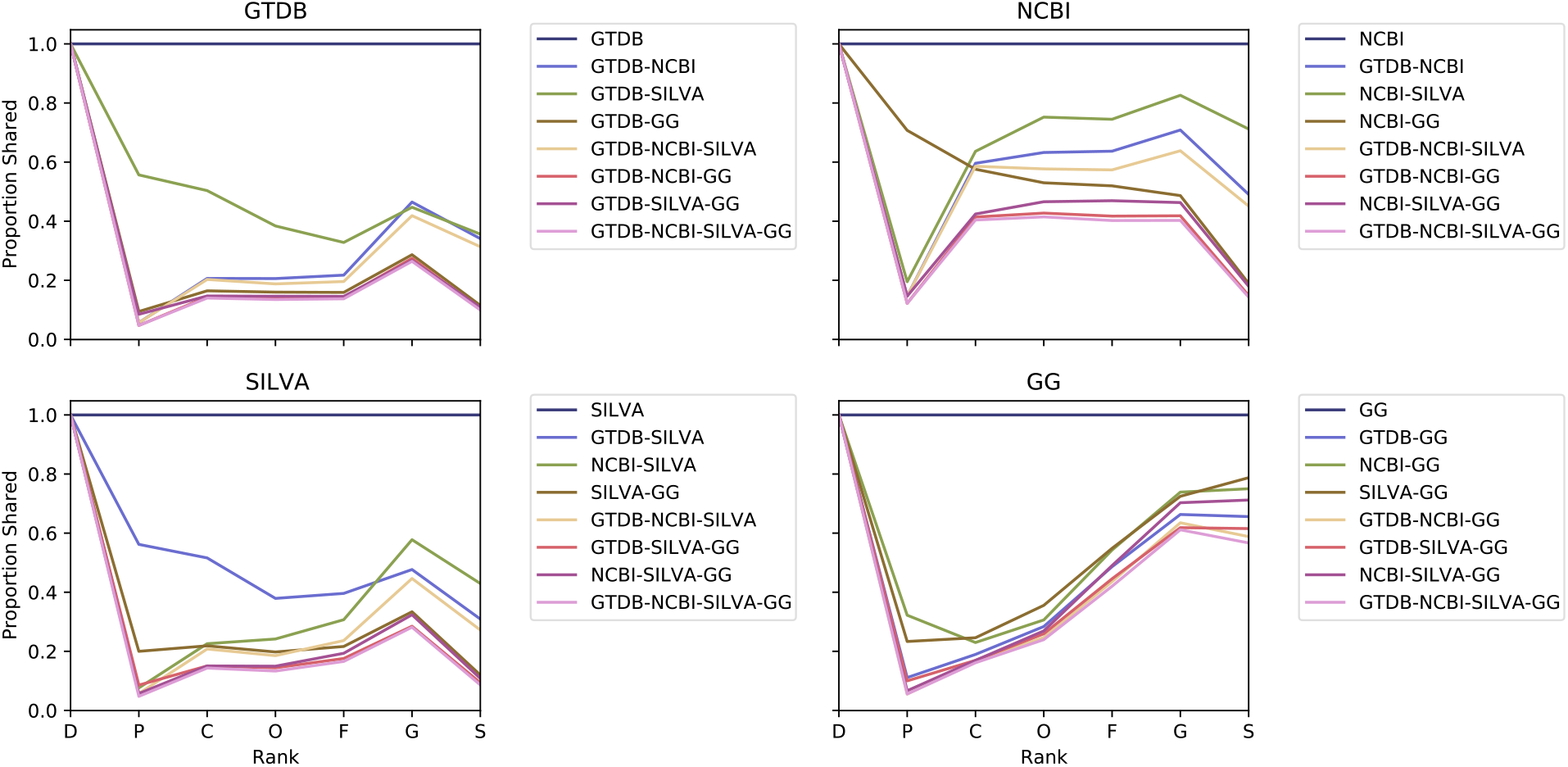
Comparison of taxonomic coverage among SILVA, Greengenes, GTDB, and NCBI-RefSeq 16S rRNA gene databases. Each panel displays the proportion of taxa represented in one reference database (as indicated in the panel title) that are shared with each other database at each taxonomic rank. The legends indicate which groups are being compared: the reference alone (always 1.0, shown for clarity), pairs consisting of the reference and one other database, trios consisting of the reference and two other databases, and the proportion of the reference’s labels that are shared by all four databases. Rank labels on x-axis: D = Domain, P = phylum, C = class, O = order, F = family, G = genus, S = species.

We found that very low proportions of taxonomic labels were shared between and among databases at all ranks below domain. In general, select pairs shared more labels than trios, and ∼30–50% (of the total number of labels in each of those databases) of species labels were shared by SILVA, GTDB, and NCBI-RefSeq (Figure 4). Proportions increased at the genus rank, and for some groups at the species rank, reflecting differences in taxonomic labels (often due to taxonomic reclassifications related to database age) at the intermediate ranks. Taxonomic reclassifications have rendered many of the older taxonomies obsolete, most notably (and unsurprisingly) the Greengenes 2013 release taxonomy, as reflected in the low proportions shared with all other databases. Proposed taxonomic reclassifications in GTDB are unique to that database, leading to reduced sharing with all other databases. SILVA exhibited relatively low proportions of genus and species labels shared with other databases, but this is unsurprising given that SILVA does not curate species labels (as discussed above). Considering these limitations, these findings indicate that a reasonably high proportion of genus and species labels are shared across databases, and instances of non-sharing reflect taxonomic reclassifications more often than lack of coverage. Greengenes exhibited notably poorer taxonomic coverage, taking into account both the paucity of unique labels (Figure 3A), low taxonomic entropy (Figure 3B), the markedly lower level of sharing with all other databases (compared to other pairs of databases), and the high level of coverage of Greengenes taxonomies by the other databases (Figure 4).

Using NCBI-RefSeq as the standard for coverage of official taxonomic names (as the only taxonomy in this comparison that is comprised mostly of genomes sequenced from type material), SILVA exhibits the best coverage at class through species rank (Figure 4), but GTDB exhibits only slightly lower coverage at those ranks (most likely because proposed reclassifications reduce the degree of taxonomic sharing), hence both exhibit similar levels of coverage. By comparison, only a small proportion of GTDB and SILVA species labels are shared with other databases, reflecting both taxonomic inconsistencies (as described above) as well as the inclusion of uncultured and proposed taxa. Taken together, these findings reinforce the suggestion that NCBI-RefSeq may contain the best coverage of accepted type strains, making it best suited to some applications, although the greater inclusion of non-type and uncultured species in SILVA and GTDB may make these databases more suitable for environmental survey applications and other studies containing many uncultured organisms.

### Effects of processing steps on the SILVA 16S rRNA gene database

RESCRIPt contains multiple functions that were designed specifically for handling data from SILVA, due to the popularity of SILVA as a reference for LSU and SSU rRNA gene sequences (Figure 1). We tested the impact of several of these steps on the SILVA 138 release 16S rRNA sequences to inform best practices for processing these data with RESCRIPt. Removing abnormally short sequences (fragments) (Figure 5A) and sequences with excessive ambiguity and homopolymer content had the beneficial effect of reducing database size (Figure 5B) without substantial loss of sequence entropy (Figure 5C). Ambiguously labeled taxa (e.g., those unidentified at genus or species ranks) may create “taxonomic noise” but clearly represent unique genotypes and should not be removed (Fig 5B). Whereas the strict removal of any sequence containing ambiguous taxonomic annotations (typically at the lower ranks) resulted in the removal of 305,636 sequences. This had a noticeable effect on sequence entropy.

**Figure 5.**
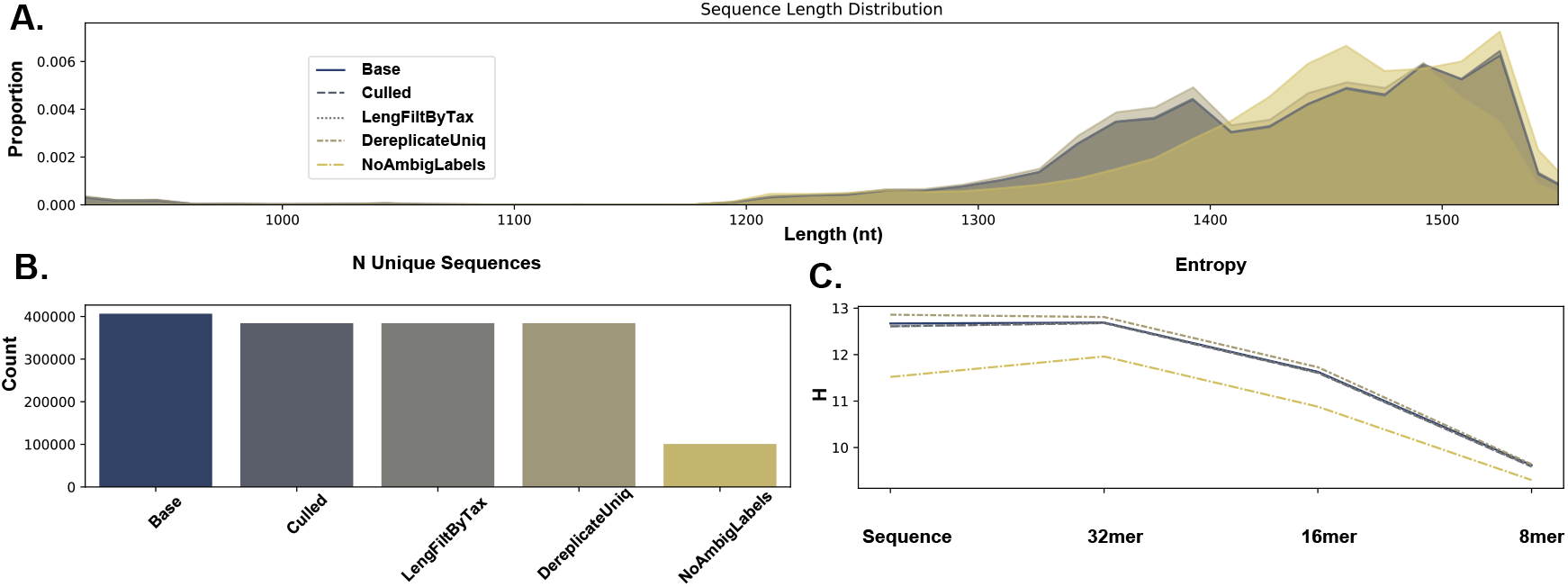
Comparison of sequence information across each successive sequence quality filtering step as applied to the SILVA 16S rRNA gene database. A, Sequence length distributions. B, Number of unique sequences. C, Entropy of full-length sequences and different kmer lengths. Note: The subsequent sequence length filtering did not have any effect on the data as the NR99 reference database is already pre-trimmed as specified above. Base: the complete NR99 SILVA database, Culled: after sequences with either 8 or more homopolymers and/or 5 ambiguous bases removed, LengFiltByTax: sequence length filtering of the data based on taxonomy, i.e. removal of Archaeal and Bacterial sequences less than 900 and 1200 bp in length, respectively. DereplicateUniq: Taxonomy and Sequence dereplication using “uniq” mode (i.e. any identical sequences with differing taxonomy will not be merged), NoAmbigLabels: any sequence data associated with ambiguous labels (typically at lower taxonomic ranks) are removed from the data set.

The evaluate-taxonomy action was also used to examine the number of unique taxonomic labels (Figure 6A), taxonomic entropy (Figure 6B). Optimal classification performance for each classifier without cross-validation (Figure 6C) as was performed earlier (Figure 3D), and with cross-validation (Figure 6D). Aside from the strict removal of ambiguous taxonomic annotations, quality filtering also had minimal impact on classifier performance and entropy,

**Figure 6.**
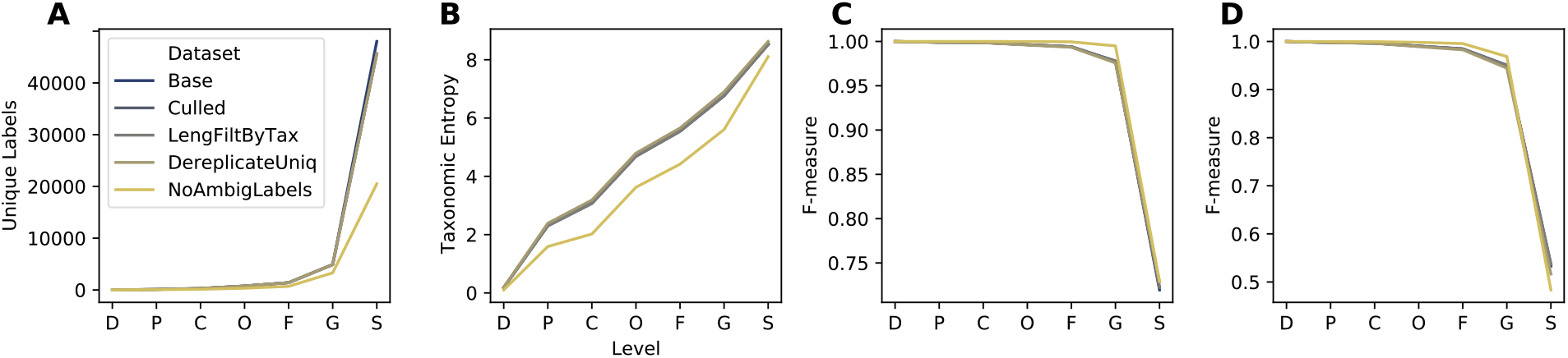
Comparison of taxonomic information and simulated classification accuracy across several successive steps of quality filtering of the NR99 16S rRNA gene databases. A, Number of unique taxonomic labels; B, Taxonomic entropy; C, optimal classification accuracy from the evaluate-fit-classifier action (as F-Measure) without cross-validation (simulating best possible classification accuracy when the true label is known but classification accuracy may be confounded by other similar hits in the database); D, optimal classification accuracy from the evaluate-cross-validate action (as F-Measure), which simulates pseudo-realistic classification task whereby a set of query sequences may not have an exact match in the reference database. See Figure 5 Legend for label descriptions. Rank labels on x-axis: D = Domain, P = phylum, C = class, O = order, F = family, G = genus, S = species.

### Effect of Clustering on Sequence and Taxonomic Information: lessons from the Greengenes 16S rRNA gene database

Clustering sequences into OTUs has long been practiced for dereplicating and reducing errors in marker-gene sequencing experiments [72]. In times of yore, clustering was also often applied to reference sequence databases for marker-gene sequencing, to reduce complexity and thus computational requirements. We have previously shown that clustering COI diet metabarcoding databases at 97% vs. 99% is detrimental to database information [73], but the effects of clustering on database quality more generally are lacking. To benchmark general effects of OTU clustering on database quality, and for 16S rRNA gene sequencing specifically, we used RESCRIPt to evaluate multiple database quality characteristics of the Greengenes database (13_8 release) [21] clustered at multiple OTU % similarity thresholds. The Greengenes public release data contain pre-clustered sequence and taxonomy files (with LCA consensus taxonomies assigned to OTU clusters), which were evaluated in this benchmark using the RESCRIPt actions evaluate-taxonomy, evaluate-fit-classifier, evaluate-cross-validate, and evaluate-classifications.

The loss of information as a result of clustering sequences has been highlighted by others for bacterial SSU [42,74] and metazoan COI [73] query sequences. We build on their work by demonstrating similar issues with the use of OTU clustering for reducing complexity in marker-gene sequence reference databases. Decreasing % similarity threshold rapidly leads to information loss; at the genus and species ranks the number of unique taxonomic labels rapidly declines (Figure 7A), as sequences (and genera and species) are collapsed into larger OTUs with less taxonomic resolution. Taxonomic entropy (Shannon’s entropy [75] applied to vectors of taxonomic label counts), which measures both the richness and evenness of taxonomic labels, registers a gradual decline as OTU % similarity is decreased from 99% to 88% at both genus and species levels, and rapidly declines thereafter (Figure 7B). This indicates that, although unique genus and species labels are being collapsed into larger family-level OTUs, OTU clustering is also initially reducing label redundancy, leading to increased evenness. The proportion of terminal labels (i.e., the rank at which taxonomic annotation terminates) illustrates how the rank assignment landscape changes: a higher proportion of genus- and species-level annotations in the 99% OTU sequences gives way to a higher proportion of class-, order-, and family-level terminal annotations as the % similarity threshold is decreased (Figure 7C). Contrary to this trend, “best-case” classification accuracy (with evaluate-fit-classifier, which trains and tests a naive Bayes taxonomy classifier on the same input data without cross-validation) is seen to increase from F=0.88 to F=1.0 as databases are clustered (Figure 7D), but this phenomenon reflects the loss of information with increased OTU clustering, suggesting that the higher apparent accuracy is actually an indicator of lost database coverage. Cross-validated classification accuracy (with evaluate-cross-validate) gives a more realistic demonstration of performance, demonstrating that classification accuracy actually declines as OTU clustering % increases, as database coverage decreases making the classifier less effective (Figure 7E). Taken together, these results suggest that even very modest OTU % clustering thresholds are likely to negatively affect database information content as well as classification accuracy. Although a small degree of dereplication and clustering may be beneficial for reducing sequence and taxonomic redundancy, we recommend against using OTU clustering < 99% similarity for any marker-gene sequence databases.

**Figure 7.**
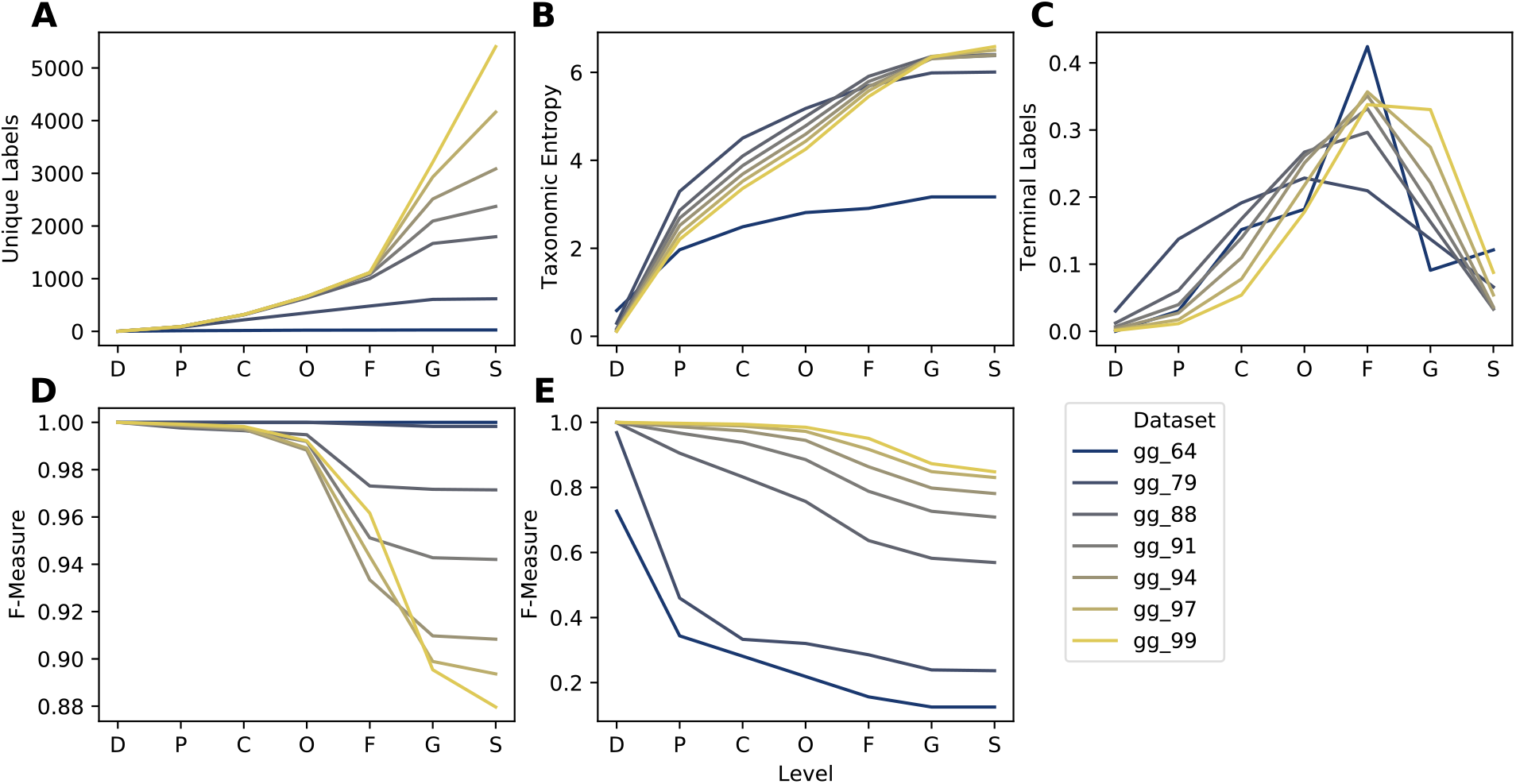
Taxonomic information (A-C) and classification accuracy (D-E) of Greengenes 16S rRNA gene database clustered at different similarity thresholds. Subpanels show taxonomic/classification characteristics at each taxonomic level: A, Number of unique taxonomic labels; B, Taxonomic entropy; C, number of taxa that terminate at that level; D, optimal classification accuracy (as F-Measure) without cross-validation (simulating best possible classification accuracy when the true label is known but classification accuracy may be confounded by other similar hits in the database); E, classification accuracy (F-Measure) with cross-validation (simulating realistic classification tasks when the correct label is unknown). Rank labels on x-axis: D = domain, P = phylum, C = class, O = order, F = family, G = genus, S = species.

### Reference Curation Improves Taxonomic Classification: lessons from the UNITE Fungal ITS database

Sequence reference databases are often subsetted by investigators to focus on particular clades of interest or to perform additional curation of public datasets. Some researchers have generated environment-specific databases, founded in the belief that such databases increase taxonomic classification accuracy by removing sequences that are genetically related but ecologically distinct from species found in a specific environment [52,55–59], although this can elevate the risk of false-positive errors [76]. RESCRIPt contains several methods to support and evaluate such filtering decisions, which then become embedded in provenance to facilitate transparent and reproducible use of these databases. To demonstrate this filtering capacity, and evaluate its effect on classification accuracy, we performed a benchmark of the UNITE [30] ITS database. The most recent releases of UNITE contain different versions that we benchmark here: (1) different “species hypothesis” OTU clustering thresholds (including a dynamically defined clustering threshold defined by manual curation [30]); (2) release versions containing ITS sequences for all eukaryotes vs. only fungi; and (3) the fungal database filtered (by RESCRIPt) to contain only sequences that are annotated at the order level or below.

OTU clustering (97% vs. 99% vs. dynamic clustering) exhibited minimal impact on results, though the 99% OTUs yielded the highest taxonomic information and classification accuracy (Figure 8), corresponding to the more comprehensive clustering benchmark performed above (Figure 7). As expected, the “all eukaryotes” version of UNITE contains more than twice as many sequences as the fungi-only database (Figure 8A), though taxonomic entropy is slightly lower (Figure 8B), reflecting a greater proportion of non-fungal sequences that are not annotated at the family, genus, and species ranks (Figure 8C). Taxonomic classification accuracy is lower in the “all eukaryote” release version of the UNITE database, compared to fungal sequences alone (Figure 8D-E). This indicates that removing non-target sequences from the database improves taxonomic resolution; however, such practices (whether focusing on particular clades or environment-specific species) is fraught with risks and should be used with caution. If the non-target sequences can be detected (e.g., amplified by the same primer, or introduced by cross-contamination or rare migration events), filtered databases may lead to misclassification (e.g. a sequence may be classified as a fungus, when it is actually a metazoan, or vice versa [76]). We recommend careful consideration of these risks when selecting primers, databases, and filtering decisions for marker-gene and metagenome sequencing studies. An advantage of using RESCRIPt for database construction and curation is that these processing steps are embedded in provenance, allowing the appropriateness of these steps to be re-evaluated at later stages (e.g., in documenting results, peer review, and re-use in future studies and by other researchers).

**Figure 8.**
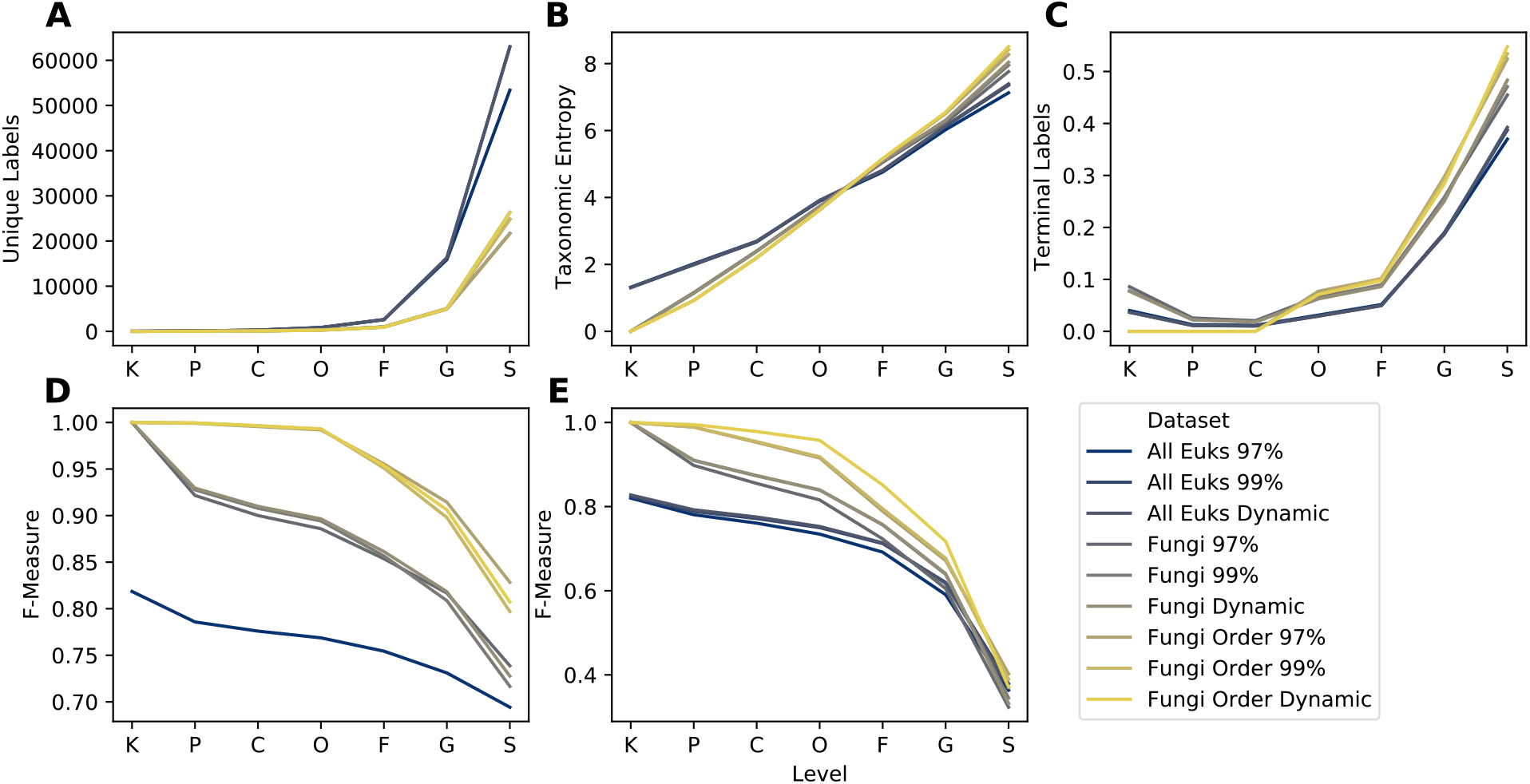
Taxonomic information (A-C) and classification accuracy (D-E) of UNITE ITS domain database with different filtering and clustering settings. Filtered versions include the “all Eukaryotes” 2020.04.02 release version containing all Eukaryotes (“All Euks”), filtered to contain only Fungi, and filtered to contain only Fungi with at least order-level taxonomic annotation (“Fungi Order”). Cluster levels indicate which UNITE release version was used: sequences clustered at 97% similarity, 99% similarity, or the UNITE “dynamic” species hypothesis threshold. Subpanels show taxonomic/classification characteristics at each taxonomic level: A, Number of unique taxonomic labels; B, Taxonomic entropy; C, proportion of taxa that terminate at that level; D, optimal classification accuracy (as F-Measure) without cross-validation (simulating best possible classification accuracy when the true label is known but classification accuracy may be confounded by other similar hits in the database); E, classification accuracy (F-Measure) with cross-validation (simulating realistic classification tasks when the correct label is unknown). Rank labels on x-axis: K = Kingdom, P = phylum, C = class, O = order, F = family, G = genus, S = species. See Materials and Methods for more details on how these databases were created and processed.

Filtering out fungal sequences that were not annotated at least to order level removed only a small fraction of unique labels (Figure 8A), boosting entropy by a narrow margin (Figure 8B) and classification accuracy by a wide margin (Figure 8D-E). These results indicate that, even in curated release versions of some public databases, some additional curation is beneficial to remove sequences with missing or uninformative taxonomic labels. Filtering with RESCRIPt enables researchers to automatically record these filtering decisions in provenance, making it clear when, where, and why their reference materials diverge from public release versions of these databases.

### Clustering and primer-region trimming effects on a BOLD COI gene database

The COI gene is a common target for taxonomic identification of metazoa, both for diet metabarcoding and eDNA studies [34]. The earliest versioned COI databases were available through a single resource: the Barcode of Life Data Systems (BOLD) [33]. However, a growing number of researchers have recently contributed updated COI databases using either BOLD or NCBI GenBank (or both) as source data [14,77–80]. We conducted a series of benchmarks that evaluated the compositional and performance effects of COI databases constructed by varying the following: first, clustering reference sequences and/or using full length sequences versus trimming references to a particular primer region; second, the reference source itself (BOLD vs. NCBI GenBank) [33,81]. The BOLD vs. NCBI GenBank comparison is the subject of the next section. For all COI benchmarks, the results are reported separately for the two largest groups of COI sequence data: arthropods and chordates.

Previous reports have separately described the reduction of taxonomic information when clustering COI sequences, as well as the effects of classifier performance based on the reference source [73,79,80]. We build on those previous works to demonstrate the effects of clustering in combination with trimming these sequences to a particular region within the COI sequence, focusing only on reference sequences obtained from BOLD. Clustering sequences dramatically reduces the number of unique sequences in both the untrimmed and primer-trimmed COI datasets (Figure 9A). Similarly, trimming COI sequences to a particular primer-defined region results in a decrease in the number of unique chordate and arthropod sequences. Nevertheless, it is worth noting that despite trimming to a region containing sequences approximately 180 bp in length, a large amount of sequence diversity remains for both arthropod (N=611,166) and chordate (N=69,924) references. Sequence entropy was similarly reduced when references were trimmed or clustered (Figure 9B).

**Figure 9.**
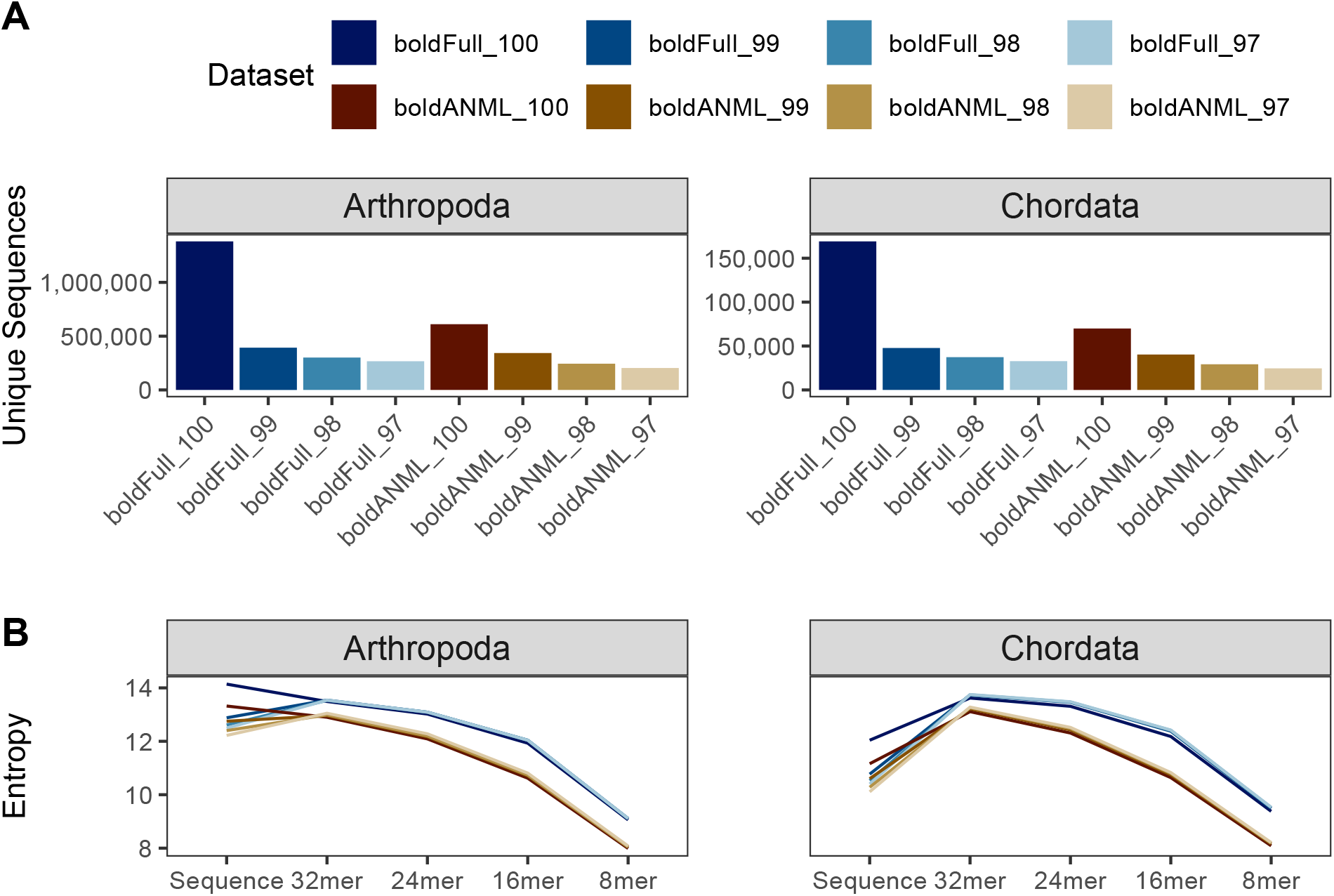
Comparison of sequence information from from BOLD COI gene database for available arthropod and chordate sequences. Differences in Dataset reflect whether sequences were trimmed to a particular primer region (boldANML) or not (boldFull), and whether sequences were dereplicated (100) or clustered at a particular percent identity (97, 98, 99). A, Number of unique sequences. B, Entropy of sequences and different kmer lengths.

We find that sequence clustering and primer trimming both also contribute to a decrease in taxonomic information, most pronounced at the species rank (Figure 10A). Untrimmed chordate sequences clustered at 97% identity (“Full_97”) contained just 77% as many unique labels as the untrimmed sequences that were only dereplicated (“Full_100”). Trimming these chordate sequences to a particular primer region also reduced taxonomic information, with dereplicated (“ANML_100”) and clustered (“ANML_97”) sequences containing 80% and 60% as many unique labels as the untrimmed dereplicated Chordate reference set, respectively. A similar effect was observed for arthropod sequences, with 97% identity clustered untrimmed sequences containing 83% as many unique labels compared to the dereplicated and untrimmed arthropod references. Dereplicated and primer-trimmed arthropod sequences, and 97% clustered primer-trimmed sequences contained 78% and 62% as many labels relative to the untrimmed, dereplicated arthropod sequences, respectively.

**Figure 10.**
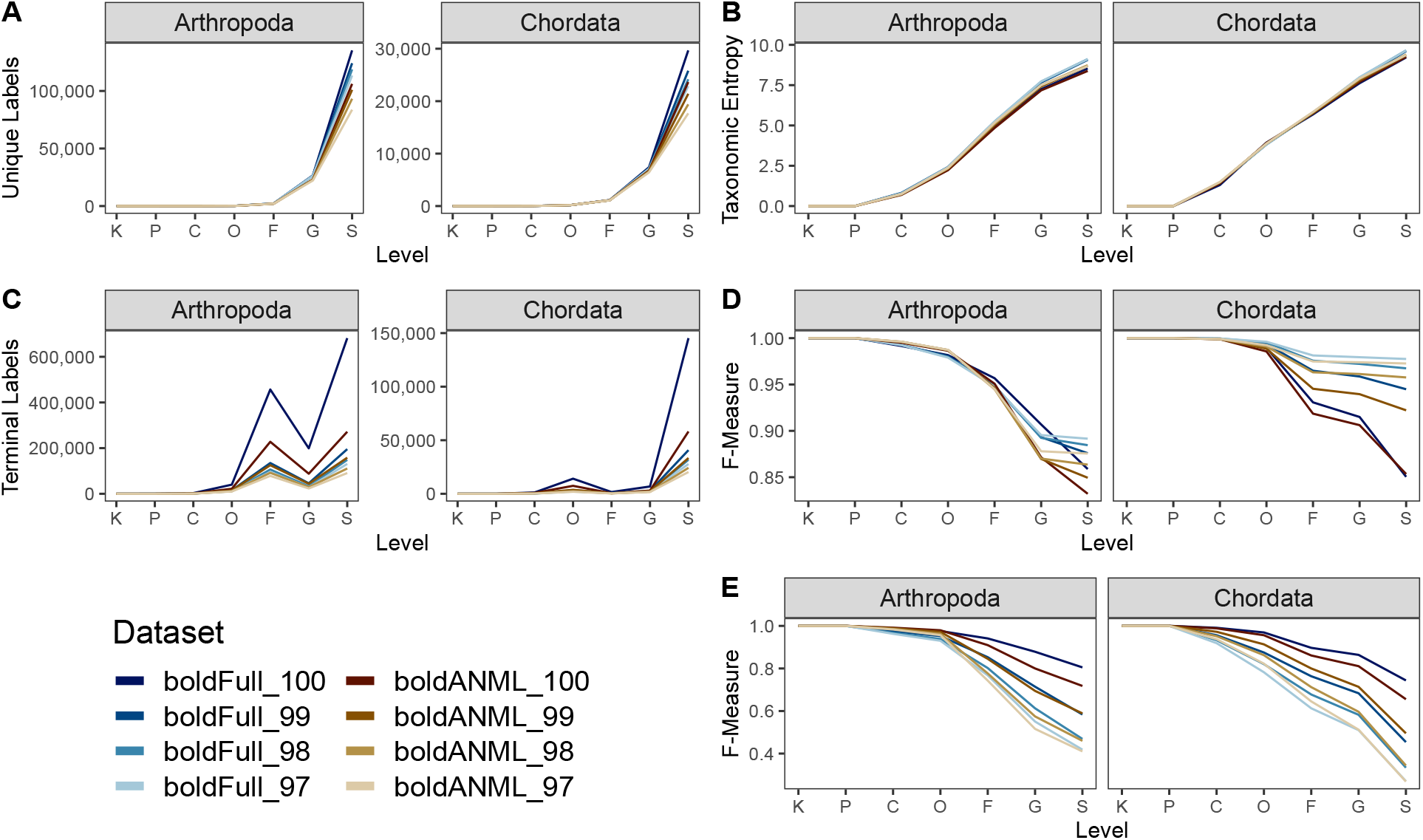
Comparison of taxonomic information and simulated classification accuracy from BOLD COI gene database for available arthropod and chordate sequences. Differences in Dataset reflect whether sequences were trimmed to a particular primer region (boldANML) or not (boldFull), and whether sequences were dereplicated (_100) or clustered at a particular percent identity (_97, _98, _99). A, Number of unique taxonomic labels; B, Taxonomic entropy; C, proportion of unclassified taxa at each rank; D, optimal classification accuracy (as F-Measure) without cross-validation (simulating best possible classification accuracy when the true label is known but classification accuracy may be confounded by other similar hits in the database). E, Classification accuracy with cross-validation. Rank labels on x-axis: K = kingdom, P = phylum, C = class, O = order, F = family, G = genus, S = species.

Unlike the observations with Greengenes SSU data, we find that the taxonomic entropy increases marginally when a larger clustering radius (decreasing percent identity) is applied (Figure 10B), indicating that clustering reduces redundancy of taxonomic labels. In addition, the same effect is observed when a reference sequence is reduced to a relatively shorter subsequence. Likewise, we observe that both clustering and primer trimming lead to a significant reduction in the number of terminal labels (Figure 10C). Although both arthropod and chordate references contain the most sequences with species-rank labels in BOLD, arthropods uniquely contain more sequences ending with family-level information than genus-level, which is perhaps an indication of the greater challenge in classifying many arthropod specimens. The entropy results, paired with the number of terminal labels at a given rank, indicate that sequence clustering and primer trimming are both more consequential with respect to reducing the total number of sequences (richness) than the redundancy of labels (evenness).

While trimming and clustering followed similar trends with respect to taxonomic information, the results of the evaluate-fit-classifier, our “best-case” classification accuracy, indicate opposing outcomes with respect to these two processes: primer trimming reduces true taxonomic classification accuracy (due to reduced sequence information and thus lowered ability to distinguish taxa), while clustering artificially increases it (by reducing genetic complexity and clustering genetically similar clades). Thus, the “best-case” accuracy for these COI sequences was obtained when references were untrimmed and clustered at 97%. Notably, the magnitude of these effects varied by taxonomic group: chordate sequences were more sensitive to clustering than arthropods, and primer trimming was more impactful for arthropods than chordates. Cross-validated classification suggests a more unified pattern with respect to classification accuracy, such that trimming and clustering both reduce accuracy. As mentioned previously with regards to clustering the Greengenes SSU OTUs, the differences between “best-case” and cross-validated accuracy are likely driven by a loss of information with increasing OTU clustering. Collectively, our data suggest that OTU clustering is detrimental for COI gene classification (Figure 10).

### Comparison of metazoan COI gene sequences in BOLD and GenBank

Next, we compared dereplicated and primer-trimmed metazoan COI reference sequences obtained from either BOLD or NCBI GenBank. Because some COI reference sequences have been deposited in both BOLD and NCBI, the NCBI data was partitioned into sequences cross-referenced to BOLD (“ncbiOB”) or not (“ncbiNB”), as well as represented in its totality (“ncbiAll”). In addition, we evaluated the number of distinct taxonomic labels shared among these databases to illustrate the degree of taxonomic information shared between groups.

Dereplicated and primer-trimmed sequences obtained from BOLD contain a slightly larger number of unique arthropod and chordate references compared to those obtained through NCBI GenBank (Figure 11A). Although many thousands of sequence accessions are cross-listed in both NCBI GenBank and BOLD, these data indicate that tens of thousands of COI sequences publicly available via BOLD are not cross-listed in NCBI GenBank. Sequence entropy was similar among all combined NCBI datasets and BOLD (Figure 11B) for both chordate and arthropod references.

**Figure 11.**
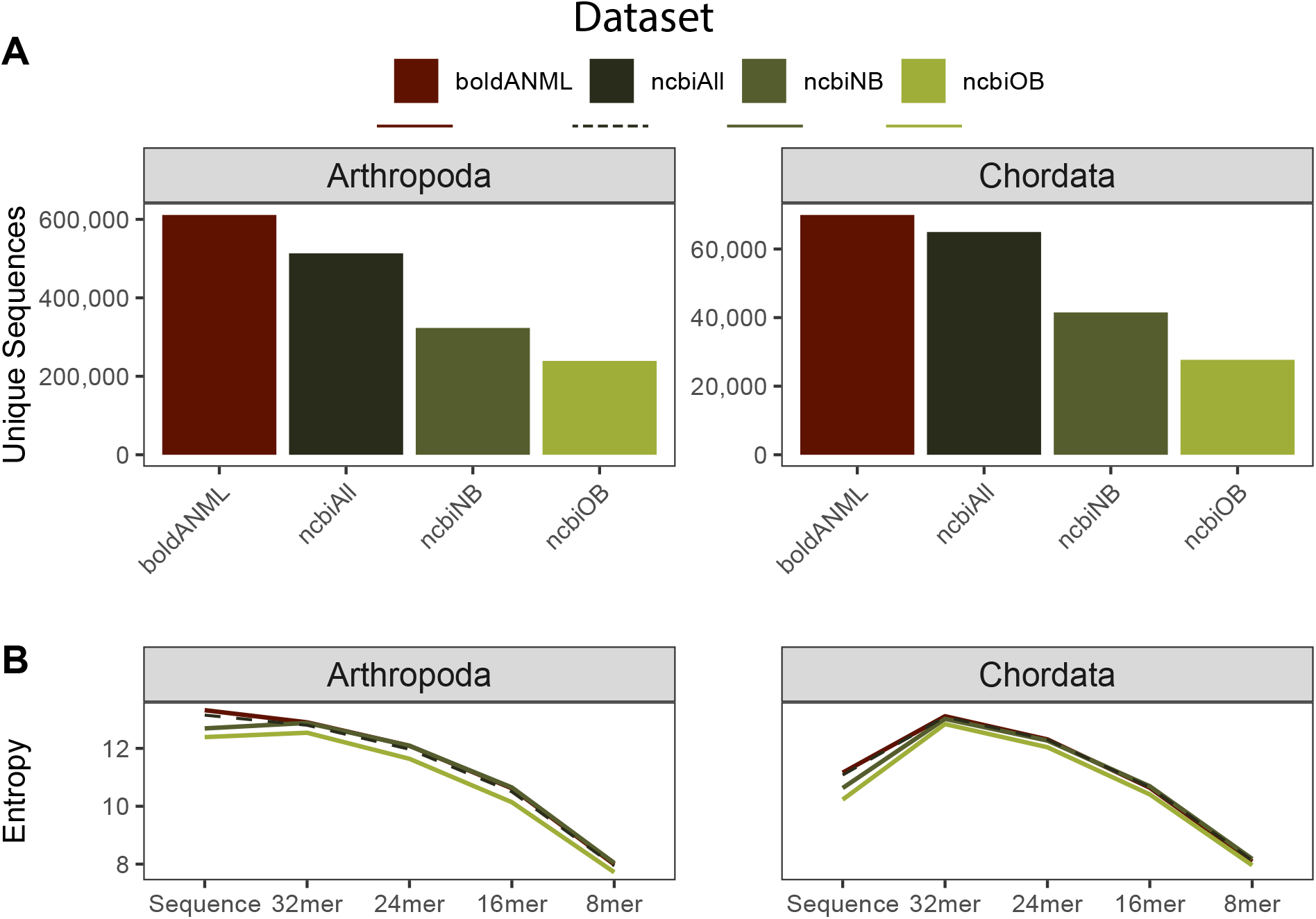
Comparison of sequence information from BOLD and NCBI GenBank COI gene databases for available arthropod and chordate sequences. All sequences were dereplicated and trimmed to a common primer region. NCBI references either contained a cross-reference term to BOLD (“ncbiOB”) or not (“ncbiNB”) or were combined together (“ncbiAll”). A, Number of unique sequences. B, Entropy of sequences and different kmer lengths.

Despite having fewer overall unique sequences, data obtained from NCBI contained more unique genus and species labels than BOLD arthropod references, and a similar number of BOLD chordate references (Figure 12A), leading to higher taxonomic entropy at the genus and species levels for NCBI (Figure 12B). Similarly, the combined NCBI database (“ncbiAll”) contains many more arthropod and slightly more chordate sequences that are assigned species labels after truncation and dereplication (Figure 12C). For arthropod references, we find that BOLD data is the least accurate database among the “best case scenario” classification (Figure 12D), but performs the best when subject to cross validation (Figure 12E). However, chordate references are consistent between both measures, and indicates that NCBI references provide an improved accuracy relative to BOLD references from family through species-levels (Figure 12D-E).

**Figure 12.**
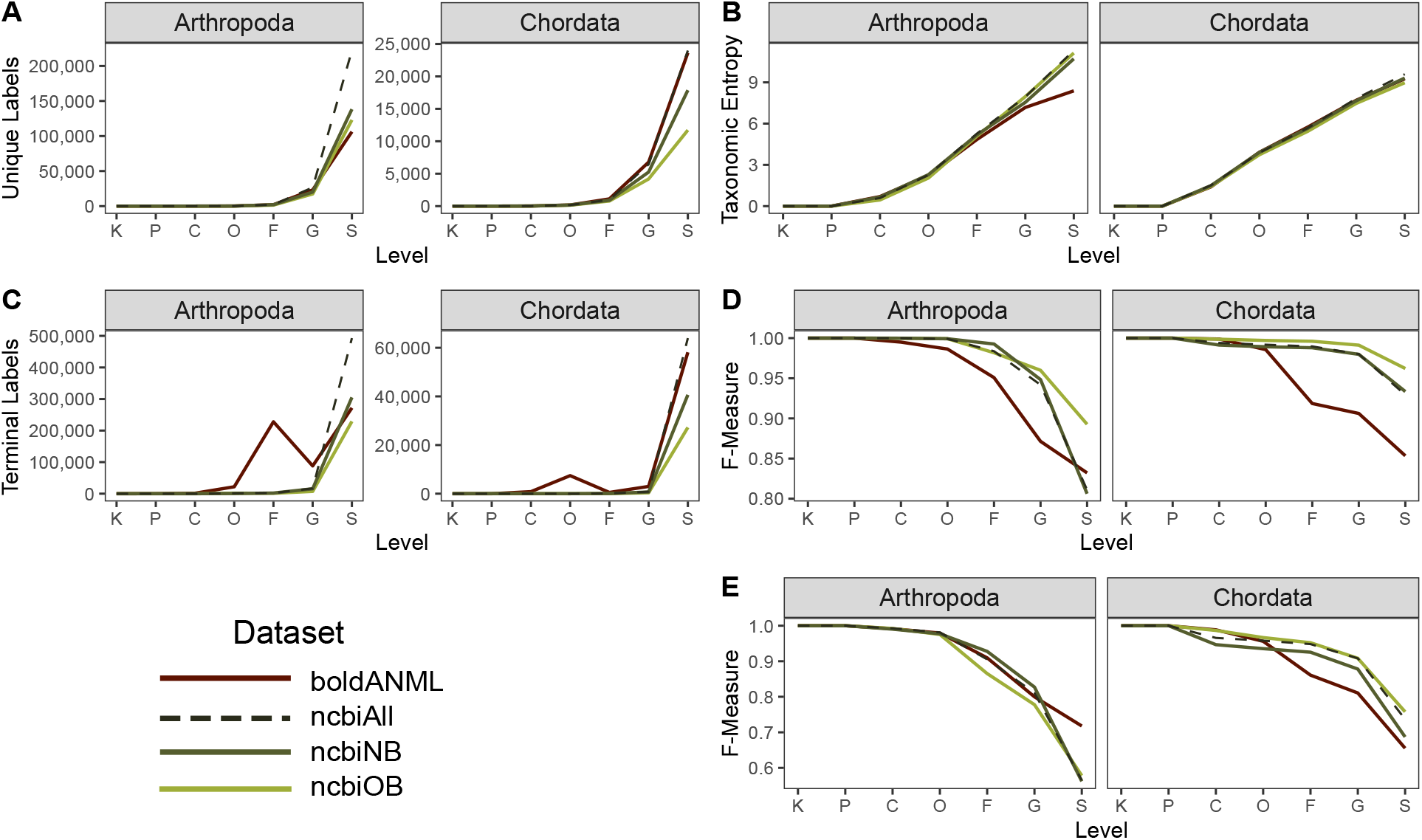
Comparison of taxonomic information and simulated classification accuracy from BOLD and NCBI GenBank COI gene databases for available arthropod and chordate sequences. All sequences were dereplicated and trimmed to a common primer region. NCBI references either contained a cross-reference term to BOLD (“ncbiOB”) or not (“ncbiNB”) or were combined together (“ncbiAll”). A, Number of unique taxonomic labels; B, Taxonomic entropy; C, proportion of unclassified taxa at each rank; D, optimal classification accuracy (as F-Measure) without cross-validation (simulating best possible classification accuracy when the true label is known but classification accuracy may be confounded by other similar hits in the database). E, Classification accuracy with cross-validation. Rank labels on x-axis: K = kingdom, P = phylum, C = class, O = order, F = family, G = genus, S = species.

### Fetching Reference Genomes for Classification

RESCRIPt’s current functionality supports retrieval of reference genomes from NCBI GenBank, enabling extensible and reproducible genomics workflows via interaction with other QIIME 2 plugins. To highlight this functionality, we used RESCRIPt along with several other plugins, q2-sourmash (https://github.com/dib-lab/q2-sourmash) [82], q2-sample-classifier [83], q2-diversity, and EMPeror [84] to generate a reproducible workflow to acquire and process a set Hepatitis E virus (HEV) genomes from NCBI-GenBank (Figure 13A), generate MASH signatures, perform pairwise genome comparisons, and visualize genome similarity via PCoA (Figure 13B). Finally, we show that HEV genotype (MASH signature) is predictive of geographic source (Accuracy = 86.1%) using k-nearest-neighbors classification with leave-one-out cross-validation (Figure 13 C). Note that this analysis was performed using a subset of HEV genomes merely as a demonstration of RESCRIPt’s broad functionality, and does not represent a fully structured test from which biological conclusions should be drawn.

**Figure 13.**
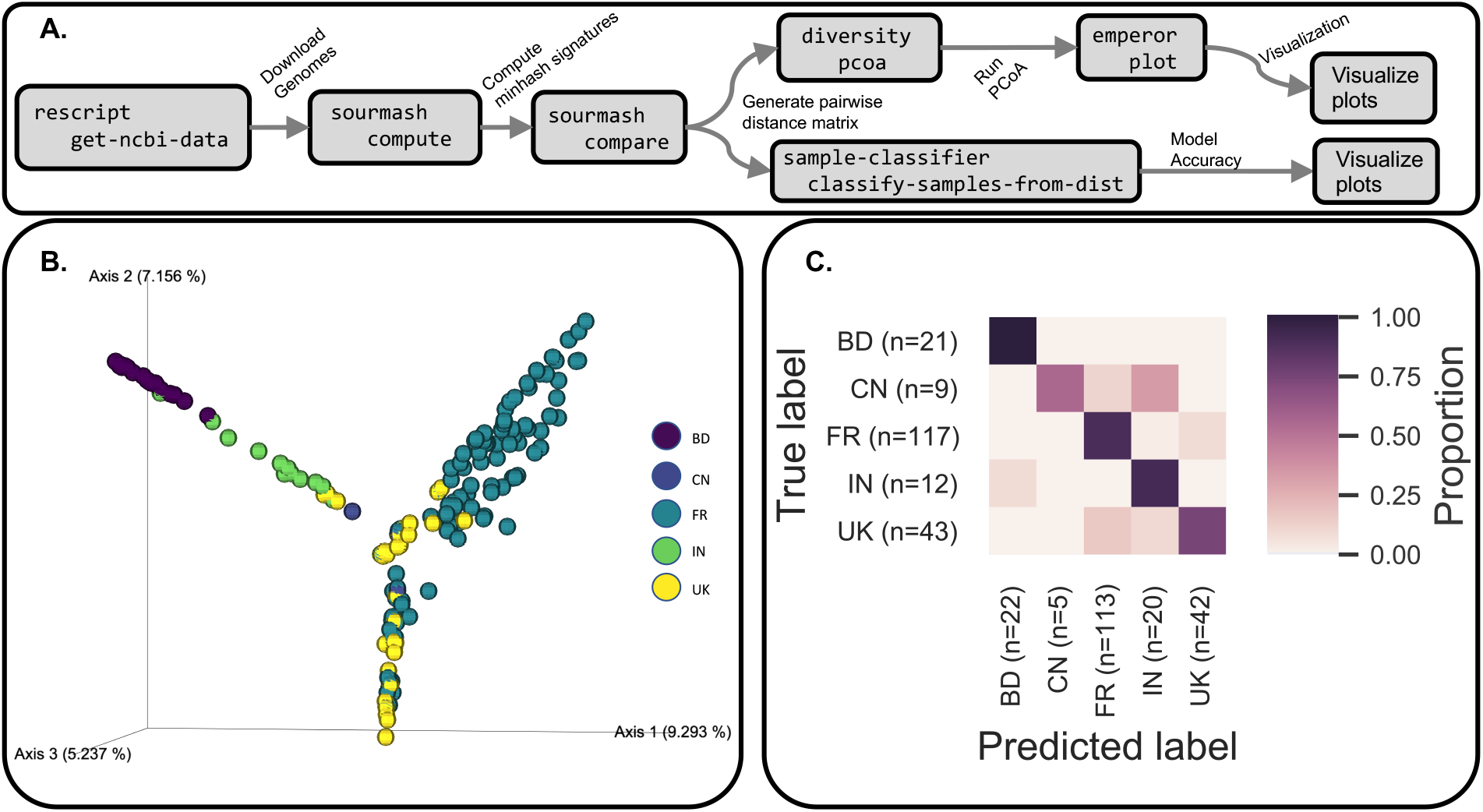
An example of using RESCRIPt for reproducible genomics workflows. HEV genomes were downloaded from NCBI-GenBank and used to make a reference genome classifier based on the following geographic locations: Bangladesh (BD), China (CN), France (FR), India (IN), and the United Kingdom (UK). The interoperability of RESCRIPt with other QIIME 2 plugins enables users to chain together a variety of functions into fully reproducible workflows that record processing decisions in data provenance. A, a data provenance graph highlighting our workflow leveraging RESCRIPt, q2-sourmash, q2-diversity, q2-sample-classifier, and EMPeror. B, PCoA plot of individual HEV genomes based on MASH signature comparison results. C, k-nearest-neighbor classification accuracy based on MASH signature dissimilarities and geographic location.

## Discussion

### The Acquisition Problem

Curated reference materials are publicly available for commonly used marker-genes such as rRNA genes [21–24,32] and the ITS region [30] for various domains of life. However, public curated reference databases are currently lacking for many other marker genes and for particular clades, creating a major bottleneck in scientific research. Even when such databases do exist, acquiring and formatting these data for use with standard methods for sequence analysis and taxonomy classification can present a steep learning curve for scientists and clinicians who lack the bioinformatics expertise required to generate and manage custom sequence and taxonomy databases. RESCRIPt resolves many of these issues, providing automatic tools for generating and formatting custom sequence and taxonomy databases (from either marker genes or genomes) from NCBI GenBank and from SILVA. Methods to provide similar support for other commonly used databases are planned for future releases of RESCRIPt. We hope that these methods will democratize the process of generating custom reference databases, supporting research efforts across the microbiome, eDNA, and metagenomics communities.

### The Reproducibility Problem

The transparent reporting, replication, and reproduction of scientific discoveries is not a new problem, but it is one that has become complicated in the digital age, as experiments and computational methods become increasingly sophisticated and datasets have become both larger and more likely to be re-used by others [63,65,66,85]. Reference database selection, and any subsequent curation, critically impact the findings from marker-gene and metagenome experiments [42,73,76,86], and hence must be carefully documented to allow others to interpret scientific findings. For example, inappropriate use of environment-specific databases have been shown to yield alarming false-positive rates in metagenomics datasets [76]. As reference data are circulated and re-used by other researchers, it is critical that database curation steps be transparently documented and transmitted so that the impact of these decisions can be evaluated downstream both by researchers re-using those datasets, as well as by the wider scientific community when interpreting results. To address this issue, RESCRIPt utilizes QIIME 2’s integrated data provenance tracking system [70] to record and store processing steps inside each individual file generated as part of a workflow. Hence, provenance can be retrieved from any terminal result file to document and replicate the entire processing chain, from data acquisition to filtering to downstream use (e.g., for taxonomic classification or sequence analysis). We believe that embedding provenance within reference database files should become a standard in the field whenever reference data are modified by a researcher or destined for re-use by others, and hope that others utilize RESCRIPt to facilitate greater transparency and reproducibility within the microbiome, eDNA, and metagenomics communities.

### The Curation problem

Our results have shown that formatting and correcting taxonomy and other metadata are critical components of reference database generation and management prior to applications for sequence classification, e.g., to standardize taxonomic ranks across entries [87]. We have already implemented functions in RESCRIPt to format the popular SILVA rRNA gene and NCBI GenBank databases, and are planning future support for parsing and editing other taxonomy formats, as well as mapping between these formats [71].

There are 4 codes of nomenclature as reviewed in [88], the International Code of Nomenclature for algae, fungi and plants (ICNafp; [89]) International Code of Nomenclature of Prokaryotes (ICNP; [90]), International Code of Zoological Nomenclature (ICZN; [91]), International Code of Virus Classification and Nomenclature (ICTV, [92]). Each of these have their own rules for taxonomic curation within their respective areas. Combining and curating taxonomic information across multiple databases can be an onerous task as new issues can arise when attempting to merge information across the respective authorities on nomenclature [88]. For example, not all taxonomic ranks are recognized, available, or even treated equally across the various databases. Valid rank fields in one database may not be recognised, or even useful, in other databases, *e*.*g*. INCafp formally recognizes the below-species ranks varieties (*varietas*) and forms (*forma*), which are not recognised within the other codes of nomenclature. Inconsistencies in taxonomic labeling, updates, and rank suffixes can cause additional incompatibilities between databases. Current efforts seek to standardize higher level suffixes of microbial nomenclature [93], may streamline bioinformatic extraction and inference of rank information.

In recent years, the explosion of high throughput sequencing technologies has allowed researchers to generate genomic data on many as yet uncultured microbial taxa. In fact, the rate at which novel genomic data can be acquired [94], and rapidly placed within a phylogenetic context [23], has surpassed our ability to appropriately resolve any conflicts with traditional Linnaean taxonomy. This has resulted in some proposals on how to taxonomically organize genomic data from uncultured microbes, and with greater emphasis on phylogenetic systematics [23,95]. For example, commonly used labels may have no officially recognized rank (e.g. Opisthokonta), but are quite informative, as they may refer to monophyletic grouping of taxa.

Sequences with missing or incomplete taxonomy/metadata and low-quality sequences are common in some databases. RESCRIPt introduces easy-to-use and user-customizable tools for detection and removing low-quality entries based on sequence filtering criteria (e.g., presence of homopolymer or ambiguous bases) or based on taxonomic information. Although some of these are trivial tasks for experienced bioinformaticists, using RESCRIPt to perform these functions results in those processing decisions being recorded automatically in the provenance stored in the downstream results files, so that this information can be recovered at any stage of downstream processing (provided that the data are maintained in a QIIME 2 archive format).

The aim of RESCRIPt is to democratize the tools for database acquisition, formatting, and curation, but RESCRIPt is not in itself a tool to automatically curate data. It is the responsibility of users to check the validity of their source data and to use those reference data and RESCRIPt appropriately. Even popular sequence taxonomy databases are prone to error [41], and issues with taxonomic naming, polyphyly, and inconsistent degrees of sequence similarity at different taxonomic ranks can complicate the use, accuracy, and contemporaneity of reference databases [96]. Defining definitions of taxonomic boundaries has historically been a challenge and can vary based on the characters used to define them, e.g. biochemical, metabolomic, ecological phenotypes, with more recent definitions, and circumscription of taxa, based upon phylogenetic relatedness and average nucleotide identity. However, these new phylogenetic approaches have resulted in either the lumping or splitting of taxa creating ever more inconsistencies between taxonomy and phylogeny [16,24,97,98].

We hope that by decreasing the number of technical hurdles involved with the generation and curation of custom databases, we will ease the point of entry, and create an interest in the taxonomic sciences among the research community. In the future, RESCRIPt could help facilitate data curation in large-scale citizen science projects in which the greater research community and general public can contribute to the growth and curation of sequence and taxonomy databases.

### The evaluation problem

After constructing a custom database comes the critical question: is the database actually useful and informative? How does it compare to other databases? User-friendly methods for sequence reference database evaluation are not currently available, making database evaluation and benchmarking a formidable challenge to the research community. RESCRIPt has implemented multiple methods for database evaluation, which generate publication-quality and interactive visualizations to allow users to explore and better interpret database quality characteristics. These involve both qualitative metrics, for evaluating sequence and taxonomy information within and between databases, as well as quantitative metrics for evaluating taxonomic classification performance of marker-gene sequences with different cross-validation schemes. Furthermore, we provide reproducible examples in the online tutorials (https://github.com/bokulich-lab/RESCRIPt) to guide users in the use and interpretation of these methods, to make these methods widely available and usable by the research community.

Using these evaluation methods, we have benchmarked performance characteristics of some of the most popular reference databases for bacterial/archaeal 16S rRNA gene, eukaryote ITS region, and animal COI gene sequences. This evaluation informs several conclusions. First, all of these databases may require some additional curation by end-users to improve fitness for certain research applications. This includes filtering low-quality sequences and annotations to improve database quality and classification accuracy, such as abnormally short sequences in the GTDB, SILVA, and NCBI-RefSeq databases. Second, we compare several of these databases side-by-side to measure relative performance metrics. We conclude that the size and taxonomic comprehensiveness of SILVA are major assets, though GTDB and NCBI-RefSeq may be more suitable for various applications that respectively require greater taxonomic and phylogenetic rigor. The use of genomes sequenced from type material provides these two databases with a robust taxonomic and phylogenetic backbone that enables users to link natural history and experimental science[88,99].

NCBI-RefSeq’s species records are extracted from data submissions to the International Nucleotide Sequence Database Collaboration (INSDC), i.e., NCBI-GenBank, the European Nucleotide Archive (ENA), and the DNA Data Bank of Japan (DDBJ). NCBI-RefSeq continually curates these species records from the primary data, often by collaborating with other groups, e.g. authorities in sequence data curation, taxonomic nomenclature, phylogenetic systematics, et cetera. Furthermore, NCBI-Taxonomy continually runs taxonomic consistency checks on assembled genomes with average nucleotide identity (ANI)[100]. These curational efforts result in a well integrated suite of biological information that can be interrogated through a variety of means and data types[25,88]. The GTDB extracts and curates data from both NCBI-RefSeq and NCBI-GenBank to generate a phylogeny of archaea and bacteria from roughly 120 ubiquitous single-copy proteins [23,87]. This phylogeny is used to inform microbial taxonomy, especially in cases where a given taxonomy is observed to be polyphyletic. In this case, a conservative approach is used to remove polyphyletic groups and normalize taxonomic ranks according to their relative evolutionary divergence. Any remaining polyphyletic groups are then flagged as a “regions of instability” in the hopes that future in-depth analyses will result in a stable set of classifiable taxa [23,87]. The efforts by NCBI and GTDB enables researchers to not only more accurately classify uncultured microbes, but also place them into ecological and evolutionary context based on their nearest phylogenetic and taxonomic neighbors.

The design, curation decisions, and ultimate quality of these databases must be considered carefully when applying them for particular purposes. For example, the SILVA database is not curated at species level, though the “organism name” provided in the source NCBI data are provided. RESCRIPt’s “get-silva-data” method for acquiring and formatting SILVA data can be configured to either report these organism names as the species labels in the output hierarchical taxonomy annotations, or to only report the SILVA taxonomy, which is curated from domain to genus rank. In our evaluation, we found that although SILVA species labels can be informative, 72% consist of unidentified, uncultured, or unknown organisms, and 2.5% do not match the genus. Downstream users should be aware of such caveats when using SILVA, or any reference data, and understand the limitations that these impose when interpreting results.

### What RESCRIPt does not do, and other limitations

RESCRIPt is designed to give researchers access to tools for reproducible nucleotide sequence and taxonomy database generation, curation, and evaluation. RESCRIPt is not in itself a data source or an authority on taxonomy, systematics, or data quality, and the qualitative and quantitative metrics that RESCRIPt can generate are not infallible indicators of quality or accuracy. As with any bioinformatics methods, the quality of RESCRIPt’s outputs is dependent on the quality of its inputs and the processing decisions made by the user. In general, users should use multiple metrics to guide their interpretations of RESCRIPt’s results, but also need to be aware of the composition of input data before making conclusions about database quality.

For example, classification accuracy metrics output by RESCRIPt can be artificially high if the input database is of low quality. This is clearly seen in the OTU clustering benchmark performed using the Greengenes database (Figure 7D-E): the “evaluate-fit-classifier” method, which classifies sequences without cross-validation, reports perfect and near-perfect species-level classification accuracy on the highly clustered sequences (e.g., 64 and 79% OTUs) (Figure 7D). This is, however, because these sequences are clustered to the extent that the remaining taxonomic coverage becomes relatively poor and sparse as near-neighbors become clustered into fewer OTUs (Figure 7A). This sparsity becomes reflected in the poor classification accuracy when cross-validation is used, as the lack of near neighbors leads to high misclassification rates for the 64% and 79% OTUs (Figure 7E). Using multiple classification evaluation methods and both qualitative and quantitative metrics (e.g., comparing classification accuracy to taxonomic and sequence entropy and coverage information) will help guide more robust conclusions about database quality.

### Future goals

RESCRIPt currently contains a range of tools for sequence reference database acquisition, management, and evaluation. Curation tools remain manual and qualitative. In the future, we plan to explore and incorporate methods for quantitative curation of sequences and taxonomies. For example, the methods used by GTDB [23] to inform species clusters based on average nucleotide identity (ANI) [101] could be incorporated for similar curation of species clusters in RESCRIPt. Other methods to detect and re-annotate mis-annotated and unannotated sequences would be valuable for guiding sequence curation efforts. However, methods such as these can be difficult to apply generally, and while ANI is useful for defining species clusters based on whole-genome sequences, it may not scale appropriately to incomplete genomes or marker-gene sequences [102].

Although the scope and benchmarks included in this study focus on marker-gene sequencing applications, genome and metagenome databases are already compatible with RESCRIPt. For example, the “get-ncbi-data” method could be used to automatically download reference genomes from GenBank, and the filtering and taxonomy functions are general purpose. More genome- and metagenome-focused functionality is planned for future releases of RESCRIPt, such as ANI [101] and MASH [103] for (meta)genome distance estimation, and methods for estimating the taxonomic classification accuracy of (meta)genome databases.

RESCRIPt’s developers remain committed to working with researchers to provide access to leading reference materials and reproducible and transparent reference sequence processing workflows. We plan to add more methods for sequence and taxonomy acquisition from public online databases that are commonly used by the marker-gene and metagenome research community, and welcome collaboration with database curators who want to better integrate their databases with RESCRIPt. The most up-to-date information related to feature requests, usage, and troubleshooting can be found on the project’s GitHub page (https://github.com/bokulich-lab/RESCRIPt).

## Conclusions

Generating and managing sequence and taxonomy reference data presents a bottleneck to many researchers, whether they are generating custom databases or attempting to format existing, curated reference databases for use with standard sequence analysis tools. Evaluating database quality and choosing the “best” database (when multiple competing options exist, such as for 16S rRNA genes) can be an equally formidable challenge. We developed RESCRIPt to alleviate this bottleneck, supporting reproducible, streamlined generation, curation, and evaluation of reference sequence databases. RESCRIPt uses QIIME 2 artifact file formats, which store all processing steps as data provenance within each file, allowing researchers to retrace the computational steps used to generate any given file. We used RESCRIPt to benchmark several commonly used marker-gene sequence databases for 16S rRNA genes, ITS, and COI sequences, demonstrating both the utility of RESCRIPt to streamline use of these databases, but also to evaluate several qualitative and quantitative characteristics of each database. We show that larger databases are not always best, and curation steps to reduce redundancy and filter out noisy sequences may be beneficial for some applications. We anticipate that RESCRIPt will streamline the use, management, and evaluation/selection of reference database materials for microbiomics, diet metabarcoding, eDNA, and other diverse applications.

## Methods

### Implementation

RESCRIPt (https://github.com/bokulich-lab/RESCRIPt) was motivated by the need for scientists to transparently and reproducibly generate and curate sequence reference databases. RESCRIPt is implemented as a free, open-source QIIME 2 [70] plugin, in order to leverage QIIME 2’s data provenance tracking system to ensure that users can trace the steps used to make their custom reference databases. QIIME 2 “Artifacts” (results files) consist of zip archives containing the result file (in typical, interoperable formats, e.g., FASTA for nucleotide sequence data) as well as data provenance information and other file metadata. Users unfamiliar or unwilling to work with QIIME 2 Artifacts can extract the data using either QIIME 2 or standard methods (e.g., the UnZip utility (http://infozip.sourceforge.net/) that is included in most Linux and Unix distributions), making QIIME 2 Artifacts a fully interoperable, portable solution for storing reference databases with integrated provenance information.

RESCRIPt is written primarily in Python 3 and depends on pandas [104,105] for dataframe operations; VSEARCH [106] and scikit-bio (scikit-bio.org) for parsing nucleotide sequences; numpy [107], scipy [108], and scikit-learn [109] for numerical and statistical operations; xmltodict (https://github.com/martinblech/xmltodict) for parsing XML; urllib and requests (https://github.com/psf/requests) for HTTP requests; q2-feature-classifier [110] for taxonomic classification; and matplotlib [111], seaborn [112], VEGA [113], and q2-longitudinal [114] for plotting and data visualization, including interactive visualizations.

The current release of RESCRIPt (2020.11) contains a variety of functions for retrieving, managing, and evaluating sequence and taxonomy reference databases (Figure 1). Details on specific functions, usage, and tutorials can be found at the project website (https://github.com/bokulich-lab/RESCRIPt), and are described in the sections below.

### SILVA data retrieval and taxonomy formatting

RESCIPt supports retrieval of SSU and LSU marker-gene data from SILVA via an automated method, “get-silva-data”, or manual import of the necessary sequence and taxonomy files (Figure 1). RESCRIPt parses the SILVA taxonomy, using three files as input:

- taxonomy rank (taxrank) file, containing both the taxonomic rank and taxonomy for each numeric taxonomy identifier (taxid);
- taxonomy mapping (taxmap) file, which maps each sequence accession to a taxid and the “organism name” provided by NCBI;
- taxonomy tree (taxtree) file, which contains the hierarchical taxonomy in newick format, with the taxids used as node labels, in which the daughter nodes contain the taxids of lower-level taxonomies.

The “parse-silva-taxonomy” method utilizes the taxrank, taxmap, and taxtree files to generate a consistent user-defined rank-associated taxonomy. Although the set of ranks can be configured by the user, the following ranks are extracted by default: domain (d_), phylum (p_), class (c_), order (o_), family (f_), and genus (g_). Any ranks not associated with taxonomy have their upper-level taxonomy propagated downward (i.e. the values forward filled with the last observed taxonomic value) towards lower-level ranks. This ensures general compatibility with downstream taxonomy classification tools, many of which may require non-empty fields at each rank. Rank propagation can be optionally disabled.

Finally, the user can choose to append the organism name (from the taxmap file) for use as the species (s_) rank taxonomy. We generally warn against this due to the myriad of inconsistent information found within the organism name field (based on our benchmarking results described herein), but it can occasionally be useful. If the user does decide to leverage the organism name, we currently only return the first two words, to remove subspecies-level information that is often included in the given organism name and which can degrade classification accuracy (e.g., because the extra information causes that species to be interpreted as a unique label).

Rank propagation is provided to allow users to extract more taxonomic information, rather than explicitly pulling down only the ranks of interest. For example, if a user opted to download sequence data along with only the six standard taxonomic ranks (see above), they may obtain the following taxonomic output when rank propagation is not used:

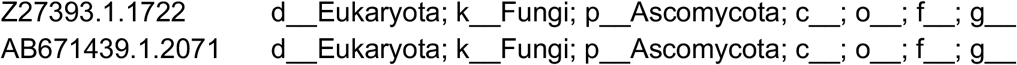

The user might assume that query sequences that “hit” either of these reference sequences would be unable to classify beyond the phylum level. However, applying rank propagation will yield the following for these same accessions:

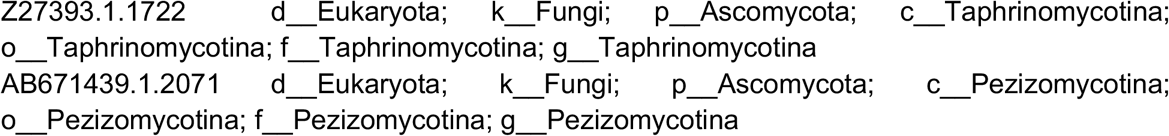

This is because intermediate ranks not selected by the user (e.g., sub-phyla Taphrinomycotina and Pezizomycotina) were propagated downward and used to fill in the unannotated ranks. Hence, forward filling allows users to disambiguate incompletely annotated reference sequences. The drawback is the conflation of taxonomy by mixing ranks from other levels.

The RESCRIPt project page (https://github.com/bokulich-lab/RESCRIPt) lists several tutorials describing how to use various RESCRIPt functions, including methods to import and parse SILVA data.

### NCBI GenBank data retrieval and taxonomy formatting

RESCRIPt supports automated retrieval of sequence taxonomy databases from the NCBI Nucleotide and Taxonomy databases [81] using the “get-ncbi-data” method. Sequences can be selected using a standard NCBI query, by specifying a list of sequence accession ids, or as a combination of the two. Downloads of large sequence databases can be made faster using parallel connections and batch downloads.

The NCBI download method retrieves the requested sequences from the NCBI Nucleotide database, cross-references their taxids with the NCBI Taxonomy database to obtain their taxonomic classifications, then standardizes the taxonomies to adhere to a fixed set of ranks. The set of ranks can be configured by the user but are kingdom (k), phylum (p_), class (c_), order (o_), family (f_), genus (g_), and species (s) by default. If a given rank is not present in the NCBI Taxonomy it is propagated down from the nearest higher rank, as described above. Rank propagation can be optionally disabled.

The RESCRIPt project page (https://github.com/bokulich-lab/RESCRIPt) lists several tutorials describing how to use various RESCRIPt functions, including methods to download and save NCBI data.

## Reference Database Benchmarks

To demonstrate some examples of how users can use RESCRIPt to process and evaluate custom reference databases from popular source data, we used RESCRIPt to conduct several benchmarks. Tutorials demonstrating this functionality, based on some of the following benchmarks, can be found on the project website (https://github.com/bokulich-lab/RESCRIPt). Workflows and data from our benchmarks can be found at https://github.com/bokulich-lab/db-benchmarks-2020 and https://github.com/devonorourke/COIdatabases. Benchmarks were designed and executed with the following aims:

1. Demonstrate various aspects of RESCRIPt’s current functionality for retrieving and curating reference sequences.
2. Compare the information content and classification performance of four commonly used sequence databases for 16S rRNA gene classification of Bacteria and Archaea, retrieved and formatted using RESCRIPt: SILVA [22], Greengenes [21], NCBI-RefSeq [25,115,116], and GTDB [24].
3. Evaluate the effects of sequence filtering on classification accuracy and information content of the SILVA [22] rRNA gene sequence database.
4. Evaluate the effects of sequence clustering on sequence and taxonomic information content, using the Greengenes [21] 16S rRNA gene sequence database.
5. Evaluate the effects of sequence filtering on classification accuracy and information content of the UNITE [30] ITS sequence database.
6. Evaluate the effects of sequence filtering on classification accuracy and information content of the BOLD [33] and NCBI GenBank [62,116] COI gene sequence databases.
7. Evaluate the effects of sequence clustering and primer-coordinate trimming on classification accuracy and information content of the BOLD gene sequence database.

Data were retrieved either using RESCRIPt (for SILVA and NCBI data) or by direct download of release data (for UNITE, Greengenes, and GTDB) or by direct download (for BOLD data; accessed July 1, 2020 and updated August 8, 2020).

SILVA data were filtered to remove sequences containing homopolymer lengths > 8 and/or > 5 ambiguous characters, using the RESCRIPt action cull-seqs with default settings. SILVA, GTDB, and NCBI data were all found to contain unusually long and short 16S rRNA gene sequences (using the RESCRIPt action evaluate-seqs), thus Archaea sequences < 900 nt [48] and Bacteria < 1200 nt [24], as performed for the SILVA releases (e.g. https://www.arb-silva.de/documentation/release-138/) were filtered out using the RESCRIPt action filter-seqs-length-by-taxon.

The raw NCBI COI data were obtained with the ‘get-ncbi-data’ action in RESCRIPt (see tutorials at https://github.com/bokulich-lab/RESCRIPt). Sequences containing the “BARCODE” keyword were considered cross-referenced NCBI data from BOLD, and labeled as “NCBIob”, while those sequences without this keyword were labeled as “NCIBnb”. These data were processed only in one fashion: initially dereplicated and primer trimmed to the ANML primer coordinates, then dereplicated once again. This produced a pair of NCBI COI databases with (“ncbiOB”) or without (“ncbiNB”) the BOLD cross-referenced label. These data were also combined, and again dereplicated, to represent the full NCBI set of COI sequences (“ncbiAll”). In all cases, every database was filtered prior to running benchmark tests to retain only those taxonomy labels and sequences associated with the phylum “Chordata” or “Arthropoda”.

The raw BOLD COI data were obtained using a custom R script (https://osf.io/m5cgs/). All data sets were filtered with similar RESCRIPt actions to remove sequences containing homopolymer lengths > 12 and/or > 5 ambiguous characters (via ‘cull-seqs’), and to retain sequences between 250 to 1600 bp (‘filter-seqs-length’). Sequences were then either dereplicated (‘dereplicate’) or clustered using one of three percent identities (97, 98, 99). These “boldFull” datasets contained sequences of variable lengths of COI sequence, and were further trimmed to the boundaries that map to ANML [117] sequences—a primer pair commonly used in animal diet metabarcoding experiments. Trimming was performed using MAFFT [118] in three stages: first, a subset of reference sequences created a high quality alignment with ‘mafft --auto’; second, primers were aligned to this small alignment file with ‘mafft --multipair --addfragments’; third, the remaining reference sequences were aligned with ‘mafft --auto --addfull’. These trimmed data were dereplicated once more to produce the “boldANML” datasets.

In several benchmarks, reference sequences and taxonomy were dereplicated and/or clustered using the RESCRIPt action ‘dereplicate’. This action uses VSEARCH to dereplicate sequences and optionally cluster them at a specified % similarity to form operational taxonomic units (OTUs), then RESCRIPt finds the most appropriate taxonomic label for each sequence cluster using one of several available modes of operation to find the last common ancestor (LCA) for the cluster, or to preserve identical sequences with unique taxonomic labels.

Taxonomic information in each database was evaluated using the RESCRIPt action evaluate-taxonomy. This action measures the number of unique labels, label entropy, and the number of unknown/unclassified labels at each taxonomic rank. Shannon’s entropy (H) [75] is defined as :

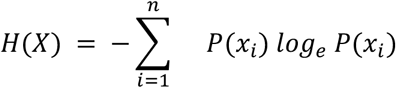

Thus, entropy relates to both the evenness and richness of information content: e.g., the number and evenness of taxonomic label frequencies or sequence/kmer frequencies.

Sequence information in each database was evaluated using the RESCRIPt action ‘evaluate-seqs’. This action measures the number of unique sequences, sequence entropy, sequence length distribution, and kmer entropy.

Taxonomic classification accuracy was simulated for each database using the RESCRIPt actions ‘evaluate-cross-validate’ and ‘evaluate-fit-classifier’, followed by accuracy evaluation with evaluate-classifications (which measures precision, recall, and F-measure in taxonomic classification results [110] and visualizes these metrics at each taxonomic rank). RESCRIPt implements two different classification evaluation methods to simulate different classification conditions, including both “unrealistic” (easy) and “realistic” classification tasks, i.e., that many of the sequences detected in a true environmental survey, e.g., of microbial or eDNA sequences in most sample types, will not have exact matches in any reference database, either because they represent novel strains, species, or higher-order taxonomic clades. The ‘evaluate-cross-validate’ action uses k-fold cross-validation (implemented in scikit-learn [109]) to perform a pseudo-realistic classification task whereby a set of query sequences may not have an exact match in the reference database, but other similar taxonomic groups may be present, as implemented and described previously [54,110]. The action splits a database into K test sets (such that each sequence appears in a test set exactly once) and classification is performed in each fold with the remaining sequences as the training set. Splitting is stratified by taxonomic groups, so that taxonomic groups are evenly stratified across training and test sets. An “expected” taxonomy is generated by this method, in which taxonomic singletons (i.e., sequence queries that do not have a representative from the same taxonomic group in the training set) are truncated so that the expected taxonomic classification is the LCA between that taxon and its nearest taxon in the training set, as this is the “correct” answer when the true taxonomic label for that query is not “known” (i.e., absent from the training set). The ‘evaluate-fit-classifier’ action trains and tests classification on the full dataset, without cross-validation, to report the best-case performance, i.e., when each query sequence has an exact match in the reference database (and hence the correct taxonomic label is known, but other matches may also be present). In this case, data leakage (where information is shared between the test and training sets) is intentional, in order to estimate the upper bound of classification accuracy for a given database by simulating an unrealistically easy classification task (using the definition of “realistic” described above). Both of these classification evaluation methods can be adapted in RESCRIPt to simulate the expected level of challenge in a given ecosystem — e.g., for well characterized sample types and clades, ‘evaluate-fit-classifier’ method may actually represent a realistic classification scenario, and users can set different levels of K for ‘evaluate-cross-validate’ (to adjust the number of splits performed) to adjust the degree of “challenge” (i.e., lower levels of K result in larger splits and more uncertainty).

In addition to dereplicating and clustering sequences, the RESCRIPt ‘dereplicate’ action is useful as a quick assessment of taxonomic resolution, versus using the classification simulation methods described above. By dereplicating the sequences and using the ‘lca’ mode to find the LCA for each cluster, followed by using the ‘evaluate-taxonomy’ visualizer (described above) to examine the number of unclassified labels at each rank, this action allows a quick assessment of how well the different taxonomic groups contained in the database can be resolved based on sequence information alone, as we demonstrate in some of our benchmarks. The classification simulation/evaluation methods are better suited for simulating realistic classification tasks, and for estimating actual taxonomy classifier performance, but are computationally intensive and time-consuming.

## Reproducible genomics workflows

To demonstrate the ability of RESCRIPt to process and compare genome data, we used the scalable MinHash (MASH) approach [103] through q2-sourmash (https://github.com/dib-lab/q2-sourmash) [82]. In brief, MASH generates compressed sketch representations of large genome sequence sets, making large genome comparisons possible through dimensionality reduction. We queried the NCBI Virus portal [119], for the Hepatitis E Virus (HEV) on September 23rd, 2020 and downloaded accessions within the viral lineage “Hepatitis E virus taxid:12461”, that were annotated as having a nucleotide completeness status of “complete”. Only HEV data with sufficient regional representation, and associated with humans, were retained. These accessions were downloaded using RESCRIPt’s “get-ncbi-data” function, and processed using a modified version of q2-sourmash (https://github.com/mikerobeson/q2-sourmash/tree/use-fasta). The q2-sourmash functions “compute-fasta” and “compare” were used to generate MASH signatures for each genome and perform pairwise genome comparisons respectively. PCoA was performed through the QIIME 2 q2-diversity plugin and visualized with EMPeror [84]. Finally, q2-sample-classifier [83], was used to determine if MASH signatures are predictive of geographic source, based on k-nearest-neighbors classification with leave-one-out cross-validation.

## Declarations

### Availability of data and materials

Source code, installation and usage instructions, and tutorials for RESCRIPt can be found at the project page: https://github.com/bokulich-lab/RESCRIPt. Workflows and data from our benchmarks can be found at https://github.com/bokulich-lab/db-benchmarks-2020 and https://github.com/devonorourke/COIdatabases/.

### Authors’ contributions

NAB conceived, designed, and wrote the source code for RESCRIPt, together with BDK and MSR. MZ and MRD contributed to the source code and testing of RESCRIPt. NAB, MSR, and DRO performed the benchmarks and analyzed the data. The manuscript was written by NAB, MSR, DRO, JTF, and BDK, and reviewed by all other authors.

